# A high-throughput, 3D microtissue platform for multiparametric analysis of tissue remodeling

**DOI:** 10.64898/2026.07.15.738540

**Authors:** Anish Vasan, Quan B Nguyen, Emily Davis, M Çağatay Karakan, Elena Westphal, Varun Shah, Wilson Wong, Emma Lejeune, Jeroen Eyckmans

## Abstract

Extracellular matrix (ECM) remodeling and force generation are fundamental drivers of tissue morphogenesis and repair, yet scalable methods to quantitatively interrogate these dynamic mechanical processes remain limited. Here, we present a high-throughput screening platform that integrates engineered three-dimensional (3D) microtissues within a standardized 96-well format. We introduce a robust mold-casting fabrication process and a layer-by-layer surface modification strategy, that ensures long-term tissue stability and prevents detachment (95% tissue formation success; stable in culture for more than 10 days). This system enables simultaneous, longitudinal quantification of tissue closure, tissue contractility, and tissue compaction from a phase-contrast imaging modality. The computational data analysis tools that accompany this framework ensure reproducibility through deterministic computation and accelerate data extraction 80-fold relative to manual annotation. Using pharmacological compounds, we show that tissue closure dynamics, force generation, and compaction represent independent variables of ECM-driven tissue remodeling, challenging assumptions embedded in commonly used contraction-based assays. Furthermore, benchmarking against reported clinical drug responses demonstrates that the 3D platform better aligns with clinical outcomes (Kendall *τ-b* = 0.72, *p=*0.045*, n* =8/10) than a conventional two-dimensional scratch wound assay (Kendall *τ-b* = 0.52, *p=*0.25*, n* =4/10). Together, this work establishes a scalable assay for functional screening and quantitative assessment of tissue remodeling dynamics in three-dimensional systems.

## I. INTRODUCTION

Tissue morphogenesis emerges from the coordinated interplay between cellular contractile forces, extracellular matrix (ECM) assembly, and ECM remodeling, which together sculpt three-dimensional tissue architecture in development, repair, and disease.^1–3^ The ECM is not a passive scaffold, its biochemical composition and hierarchical organization regulate how cells generate force while cell-generated forces reciprocally reorganize matrix architecture through protein unfolding, matrix compaction and fiber realignment.^4–7^ As a result, ECM remodeling operates across multiple hierarchical scales, from molecular turnover of matrix components to tissue-scale reorganization of matrix architecture and mechanics.^8–11^

Although biochemical assays have provided important insight into pathways governing matrix synthesis and degradation, tissue morphogenesis and repair ultimately arise from functional remodeling processes, including tissue compaction, force generation, matrix organization, and collective tissue movements that emerge from cell-matrix interactions.^12,13^ Quantifying these tissue-scale remodeling behaviors remain a major challenge in mechanobiology because existing experimental models rarely combine dynamic measurements of functional remodeling with experimental scalability.^14,15^

A major obstacle is the persistent trade-off between physiological relevance and experimental scalability. In vivo models retain physiological context but they are low-throughput and poorly suited for controlled, multifactorial perturbation studies.^16^ Conventional in vitro assays, including widely used planar scratch-wound assays^17^ and traction-based force measurement systems^18,19^, provide greater scalability but prevent matrix remodeling and tissue compaction that are key features of 3D tissue repair. More complex 3D microtissue systems, organoid platforms, and human skin equivalents and explants have advanced the study of tissue repair mechanics ^20–24^, but many remain limited in throughput, standardization, or integration with scalable screening workflows.

Here, we present a scalable 3D microtissue platform for the quantitative analysis of functional tissue remodeling. This system enables longitudinal measurements of tissue closure, contractile force generation, and tissue compaction, within ANSI-standard 96-well plate formats. Flexible micropillar force sensors are integrated with engineered microtissues anchored through an optimized surface chemistry, while WoundCompute, an open-source automated image analysis pipeline, automatically extracts multiparametric remodeling dynamics across large experimental cohorts without manual intervention. By integrating reusable 3D-printed fabrication modules with automated analysis, this framework enables quantitative dissection of functional tissue remodeling processes that have been difficult to interrogate at scale.

To demonstrate the scalability and analytical power of this framework, we performed multiparametric screening of compounds targeting cytoskeletal contractility, matrix remodeling, and wound repair pathways. This analysis revealed that tissue closure, contractile force generation, and tissue compaction are not intrinsically coupled outputs, as often implied by conventional contraction-based assays, but instead represent differentially regulated axes of functional tissue remodeling. Comparison with conventional 2D scratch assays further showed that 3D microtissue responses more closely align with clinically observed therapeutic behaviors. Together, these results establish a scalable experimental framework for quantitatively dissecting ECM-dependent functional tissue remodeling.

## II. RESULTS

### A 96-well microtissue platform for multiparametric analysis of tissue repair dynamics

Previous work from our group demonstrated that engineered stromal microtissues repair injury through coordinated tissue contraction, extracellular matrix assembly, and matrix compaction, identifying these processes as key drivers of fibrous tissue repair.^6,7,25–27^ However, these studies were performed in low-throughput formats, which limited systematic perturbation and longitudinal, multiparametric analysis. To quantitatively dissect these remodeling processes at scale, we engineered a 96-well microtissue platform that integrates three-dimensional tissue assembly and repair with simultaneous mechanical and structural readouts (Fig. 1).

**Fig. 1.**
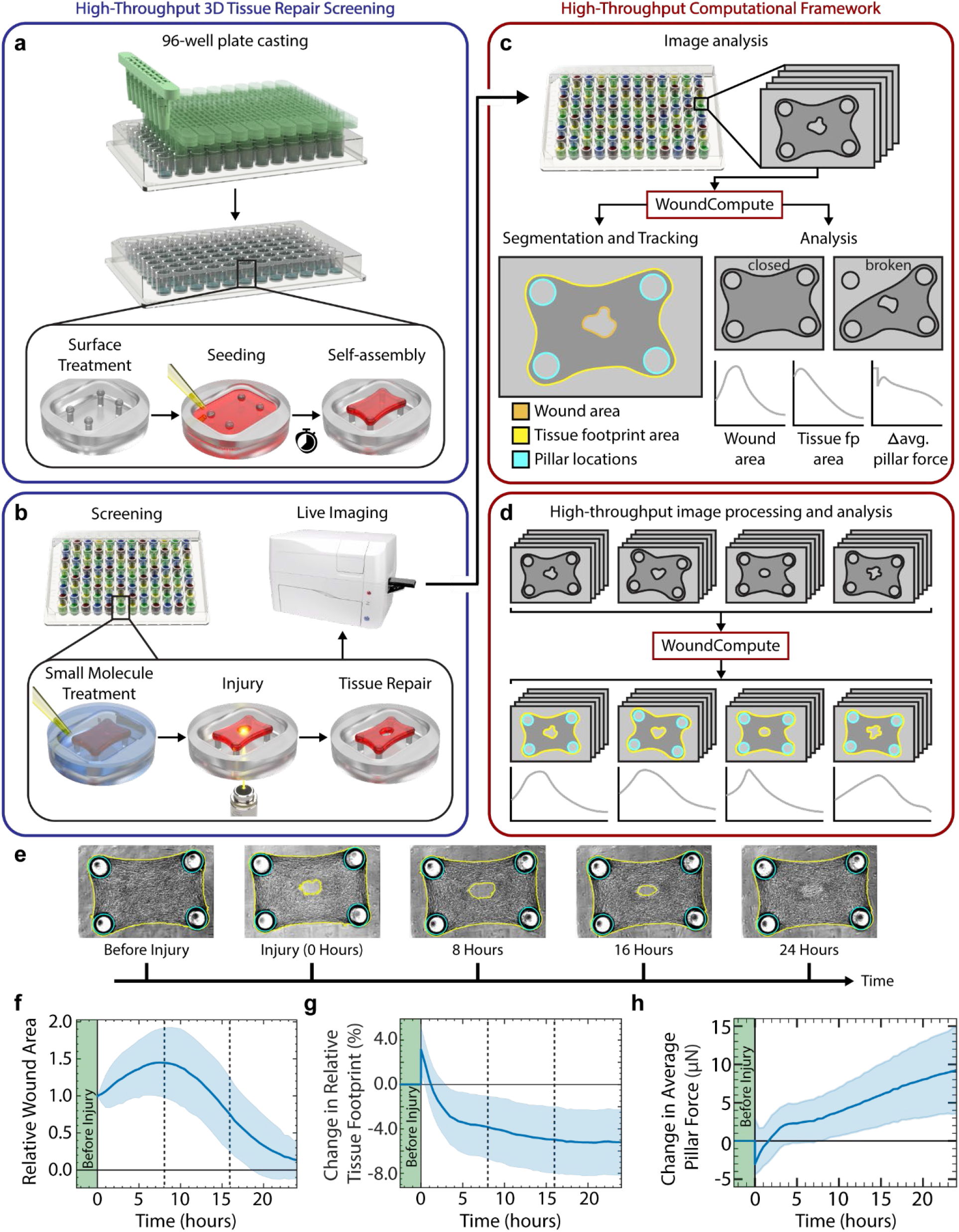
A high-throughput 3D microtissue platform to assess functional tissue remodeling. **(a)** Schematic representation of micropillar device fabrication in a 96-well plate followed by surface treatment, cell seeding and tissue self-assembly around four PDMS micropillars per well that report contractile force through their deflection. **(b)** Small molecule drug screening with tissue treatment, injury and healing process that are tracked with timelapse, phase-contrast microscopy. **(c)** Schematic representation of computational pipeline through WoundCompute being applied on images collected longitudinally from each well. WoundCompute segments the wound mask, tissue-footprint mask, and pillar locations, tracks them over time, and post-processes to classify each tissue closure and integrity. Orange outline marks the wound area. Yellow outline marks the tissue footprint area. Cyan outlines mark the pillar positions. **(d)** Schematic representation of high-throughput, parallelized processing workflow with WoundCompute where images collected from each well across the duration of the experiment are analyzed. **(e)** Representative images of an injured microtissue with cells from normal human dermal fibroblasts from neonatal foreskin tracked and segmented by WoundCompute over time. (before injury, injury at 0 h, 8 h, 16 h, 24 h) **(f)** Relative wound area (wound area normalized to the initial wound area) tracked over 24 hours (N=2, n=87). **(g)** Percent change in relative tissue footprint with respect to tissue footprint before injury tracked over 24 hours (N=2, n=86). **(h)** Change in average pillar force with respect to pillar force required at steady state before injury tracked over 24 hours (N=2, n=72). “Before injury” baseline is shaded green. (cyan = pillar markers, yellow = tissue/wound outline).

The platform combines a scalable hardware design with an integrated imaging and analysis workflow. Each well contains an array of four flexible polydimethylsiloxane (PDMS) micropillars that support microtissue formation and enable mechanical readout (Fig. 1a, b; Supplementary Fig. 1). To fabricate these devices reproducibly and at scale, we developed a reusable mold-casting system with 12 modular 3D-printed assemblies, each holding eight PDMS molds for casting individual micropillar devices directly into wells (Fig 1a, Supplementary Fig. 1a, b). After seeding normal human dermal fibroblasts (NHDFs) in a collagen type I matrix, anisotropic microtissues suspended between opposing pillars self-assembled within 12–24 hours (Fig. 1a).^25,28^ To initiate repair-like tissue remodeling, we introduced a controlled laser ablation injury and monitored the resulting tissue dynamics using time-lapse phase-contrast microscopy (Fig 1b). ^27^

The resulting image sequences were processed using WoundCompute, a computational platform developed to extract longitudinal remodeling metrics from time-lapse microscopy data (Fig 1c, d). WoundCompute segments both tissue and gap regions, enabling quantification of gap closure kinetics and tissue compaction over time. In parallel, the software tracks deflection of the spherical micropillar caps during remodeling, allowing tissue-generated forces to be calculated from established force-displacement relationships^28–30^ (Fig. 1c). Through parallelized processing, WoundCompute converts large image sequences into a scalable analysis workflow suitable for 96-well experiments (Fig. 1d). Together, this approach generates multiparametric, longitudinal datasets that capture dynamic tissue remodeling without requiring complex or multiplexed imaging modalities. (Fig1 e – h).

### Surface engineering enables scalable long-term microtissue culture

To enable long-term measurements suitable for high-throughput screening, we next sought to define the design parameters governing microtissue stability, reproducibility, and scalability. Initial implementations adapted from published methods^25–27,29–31^ exhibited higher rates of tissue malformation (15%, Supplementary Fig 2a) and detachment within 48-96 hours (Fig. 2a), limiting the utility of the 3D tissue model. We, therefore, systematically optimized two critical design parameters: initial cell seeding density and device surface chemistry. While intermediate seeding levels (4,000 cells/well) supported stable tissue formation, premature detachment persisted beyond three days with a surface coating of 2% Pluronic F-127 (Fig. 2a). To improve tissue anchorage while preserving spatial control over adhesion, we selectively functionalized spherical caps of the microtissue devices with polydopamine (PDA) to promote localized collagen and cell attachment ^32,33^, while using poly-l-lysine-grafted-polyethylene-glycol (PLL-g-PEG)^32,34^ and Pluronic F-127^30^ to passivate non-target regions. This engineered interface increased successful tissue formation to more than 95% (Supplementary Fig. 2a) and extended tissue longevity beyond ten days (Fig. 2b, Supplementary Fig. 2a, b), enabling high-throughput measurements of ECM-driven tissue remodeling over longer timescales.

**Fig. 2.**
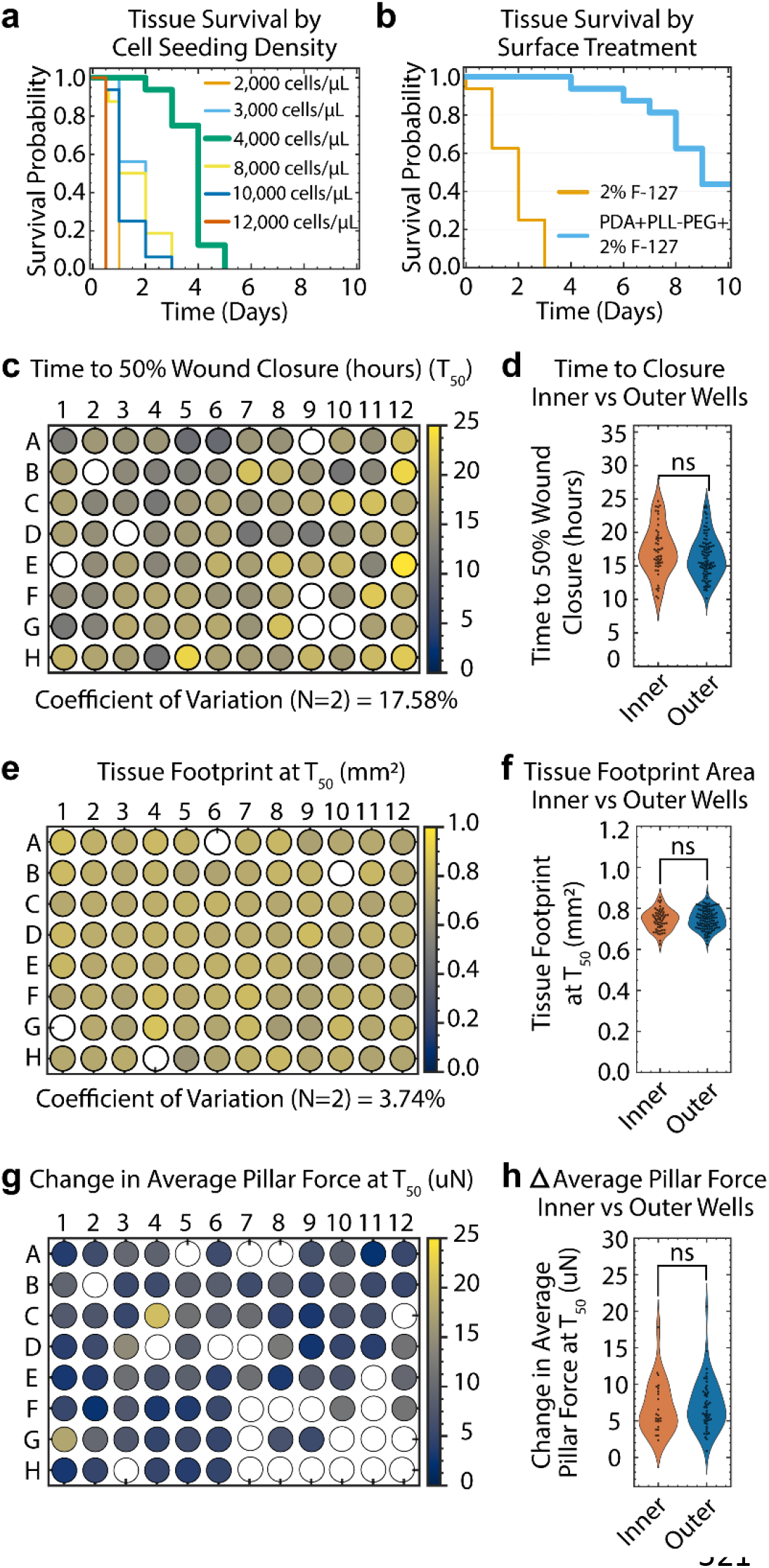
Parameter optimization for robust ECM-remodeling assays. **(a)** Kaplan-Meier plot representing tissue survival with varied cell seeding density (2000 - 12,000 cells/µL). Tissue survival is defined as the number of days a tissue remained attached to all 4 pillars. (N=1, n=12). Device surfaces were treated with 2% F-127. **(b)** Kaplan-Meier plot representing tissue survival with 2% F-127 and PDA+PLL-g-PEG+2%-F127 surface treatments to alter the cell-adhesive properties of native PDMS. Tissue survival is defined as the number of days a tissue remained attached to all 4 pillars. F-127: Pluronic F-127, PDA: Polydopamine-HCl, PLL-PEG: poly-l-lysine-g-polyethylene glycol block copolymer. (N=1, n=12) **(c)** Heatmap representing average time for each tissue to reach 50% wound area of the initial wound (T_50)_ in each well (N=2, n=87). **(d)** T_50 g_rouped by location (Outer wells: all wells located in Row A, H and Columns 1, 12; Inner wells: wells 2B to 11G). ns: not significant, unpaired t-test. (N=2, n=87) **(e)** Heatmap representing average tissue footprint in each well at T_50 (_N=2, n=86). Tissue footprint is represented from 0 to 1 mm^2^ from white to green **(f)** Tissue area at T_50 g_rouped by location (Outer wells: all wells located in Row A, H and Columns 1, 12; Inner wells: wells 2B to 11G). ns: not significant, unpaired t-test. (N=2, n=86) **(g)** Heatmap representing average pillar force generated by a tissue in each well at T_50 (_N=2, n=72). **(h)** Pillar force grouped by location (Outer wells: all wells located in Row A, H and Columns 1, 12; Inner wells: wells 2B to 11G). ns: not significant, unpaired t-test, (N=2, n=72). Empty wells are tissues/devices excluded from the analysis due to failed tissue formation, user imaging error, or 1.5 × IQR outlier.

### Platform reproducibility supports scalable microtissue screening

To support larger-scale experiments with reliable parallel measurements, we quantified well-to-well variability and spatial artifacts across the 96-well platform (Fig. 2c-h).^35,36^ Prior to injury, engineered microtissues exhibited highly uniform tissue footprint across plates, with a coefficient of variation (CV) of 7.57%, indicating consistent tissue formation (Supplementary Fig. 2c, d). After injury, the primary functional endpoint, time to 50% gap closure (T_50)_, showed moderate well-to-well variability (CV 17.58%) (Fig. 2c). Tissue footprint was highly uniform across plates (CV 3.74%) (Fig 2e), whereas the change in average pillar force showed modest variability (CV 20.47%) (Fig 3-3e). Importantly, we detected no systematic positional biases or edge effects across the plate (Fig. 2d, f, h). Together, these findings establish the platform as a reproducible and scalable assay compatible with high throughput screening (HTS) workflows ^35,36^, and show that tissue formation, closure dynamics, and contractile forces can be measured consistently across experimental replicates.

**Fig. 3.**
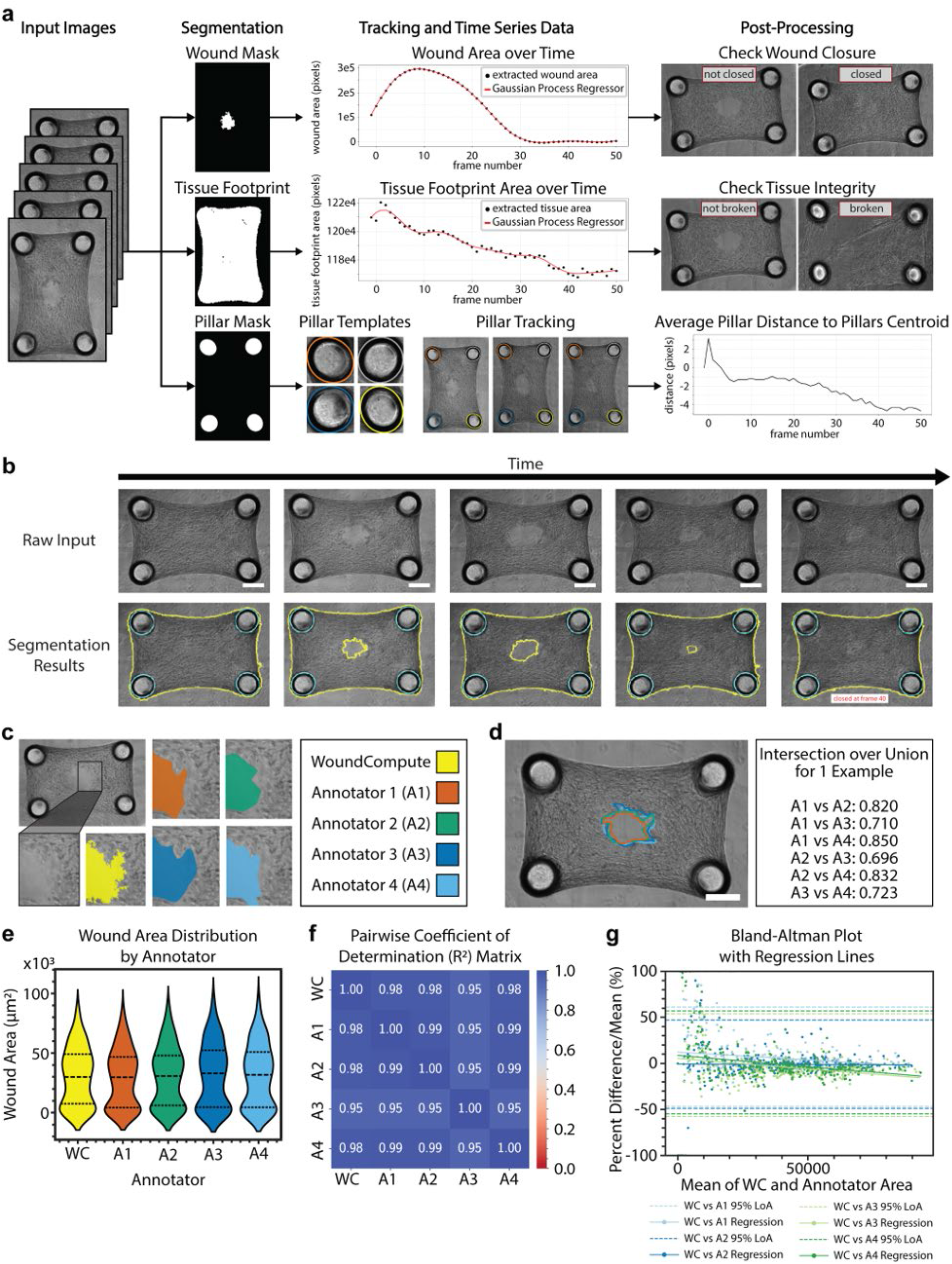
Automated extraction of tissue remodeling metrics using WoundCompute. **(a)** Schematic representation of the WoundCompute pipeline. Raw images input into the system are processed in parallel to filter and segment pillars, tissue boundaries, and wound boundaries. The package outputs the x,y positions of each pillar, wound area and tissue area. The pillar positions are used to calculate the pillar deflection and thereby calculate the tension in the tissues. **(b)** Representative images of a raw image inputs from a single tissue over time and the segmented output with pillars outlined in cyan, and yellow outline around the wound edge and tissue footprint. Scale bar: 200 µm. **(c)** Representative images of WoundCompute and manual annotators wound segmentation output. **(d)** Representative images with overlapping manual annotators wound segmentations. Intersection over union data shows percent pixel by pixel match between each manual annotator. Scale bar: 200 µm. **(e)** Absolute wound area (in µm^2^) distribution for WoundCompute and manual annotators. **(f)** Pairwise correlation matrix of wound area values from WoundCompute and manual annotators. **(g)** Bland-Altman plot comparing wound area measurements between WoundCompute and each manual annotator, with individual points for each comparison, a linear regression through these points and a dotted line representing the 95% limits of agreement.

### WoundCompute enables rapid, reproducible analysis of tissue repair dynamics

To enable analysis of large imaging datasets, we developed WoundCompute, an open-source Python-based analysis framework with graphical user interface (GUI) functionality for automated quantification of tissue repair dynamics. (see Appendix S1 for details; code available at GitHub repositories: <u>WoundCompute</u>, and <u>WoundComputeGUI</u>; image data associated with this study available at <u>10.6019/S-BIAD2719</u>). Automation was necessary due to the scale of data collected; a single 48-hour experiment with 96 wells imaged every 30 minutes produced more than 9,000 images, rendering manual annotation impractical and susceptible to user bias and annotator fatigue (see Appendix S2 for details on our Published Microscopy Dataset).

WoundCompute accepts raw phase contrast microscopy images and applies an automated segmentation pipeline to isolate the gap region, surrounding tissue, and micropillar cap positions. These segmentation masks enable quantitative extraction of closure kinetics, tissue compaction, and contractile force measurements. In addition to frame-by-frame analysis, the software performs longitudinal time-series analysis to identify complete closure events and detect structural collapse of microtissues (Fig. 3a, b).

The analysis pipeline processes individual images in approximately 12 seconds and, through parallelization, achieves an 80-fold acceleration over manual annotation. This enables full analysis of HTS experiments within approximately 3 hours 30 mins on a workstation equipped with AMD Ryzen™ 9 5900X processor (12 core), and 64 GB of DDR4 RAM running Microsoft Windows 11 (see Appendix S1 for details). By applying a fixed segmentation criterion uniformly across all samples, WoundCompute eliminates user-dependent variability and ensures greater reproducibility relative to manual annotation.

Benchmarking against independent manual annotations by multiple users revealed strong agreement between automated and manual measurements (Fig. 3c). Although most annotators achieved high segmentation overlaps, with intersection-over-union (IoU) values above 0.8, inter-user variability underscored the subjectivity of manual labeling (Fig. 3d). Pairwise coefficient of determination (R²) analysis further confirmed strong correlation between WoundCompute and manual measurement trends, indicating that the automated pipeline faithfully preserves the underlying biological trends captured by manual analysis. (Fig. 3e, f).

Notably, WoundCompute maintained segmentation fidelity during late-stage closure, a regime in which manual annotation proved less reliable. Bland-Altman analysis^37^ revealed that human annotators systematically underestimated residual gap areas in small-gap regimes, while automated segmentation maintained accuracy throughout closure progression (Fig. 3g) Collectively, these findings establish WoundCompute as a rapid, reproducible, and consistent analytical framework that complements the HTS microtissue platform.

### Multiparametric phenotyping reveals independent variables of tissue remodeling

Timelapse analysis captured a biphasic remodeling response consisting of a transient gap expansion followed by progressive, complete repair within 24 hours (Fig. 1d). These dynamics were accompanied by coordinated changes in tissue compaction, as the projected tissue footprint decreased during early remodeling before stabilizing as closure progressed (Fig. 1e). In parallel, micropillar deflection measurement revealed dynamic modulation of contractile forces characterized by an initial loss of tension immediately post-injury, followed by force recovery during closure that transiently overshot pre-injury baseline levels (Fig. 1f).

To determine whether this multivariable remodeling phenotype could resolve differential biological responses to perturbations, we next performed a pharmacological screening of 27 compounds targeting cytoskeletal contractility, growth factor signaling, integrin-mediated adhesion, and matrix degradation pathways in both neonatal and adult human dermal fibroblasts. Hits were identified using a dual-cutoff threshold based on the strictly standardized mean deviation (SSMD) of the time to 50% gap closure (*T_50_*_)_ and the relative gap area.^38^ The screen revealed pronounced asymmetry in remodeling responses. PDGF-BB was the sole compound that accelerated tissue closure, consistent with clinical reports and animal studies.^39,40^ In contrast, eight compounds significantly delayed closure, separating into four mechanistic categories targeting tyrosine kinase signaling (Dasatinib, Nintedanib)^41^, cytoskeletal contractility (Y-27632, Blebbistatin)^42^, integrin-mediated adhesion (Cilengitide)^43^, and matrix metalloproteinase inhibitors (Marimastat)^44^. (Fig 4a)

**Fig. 4.**
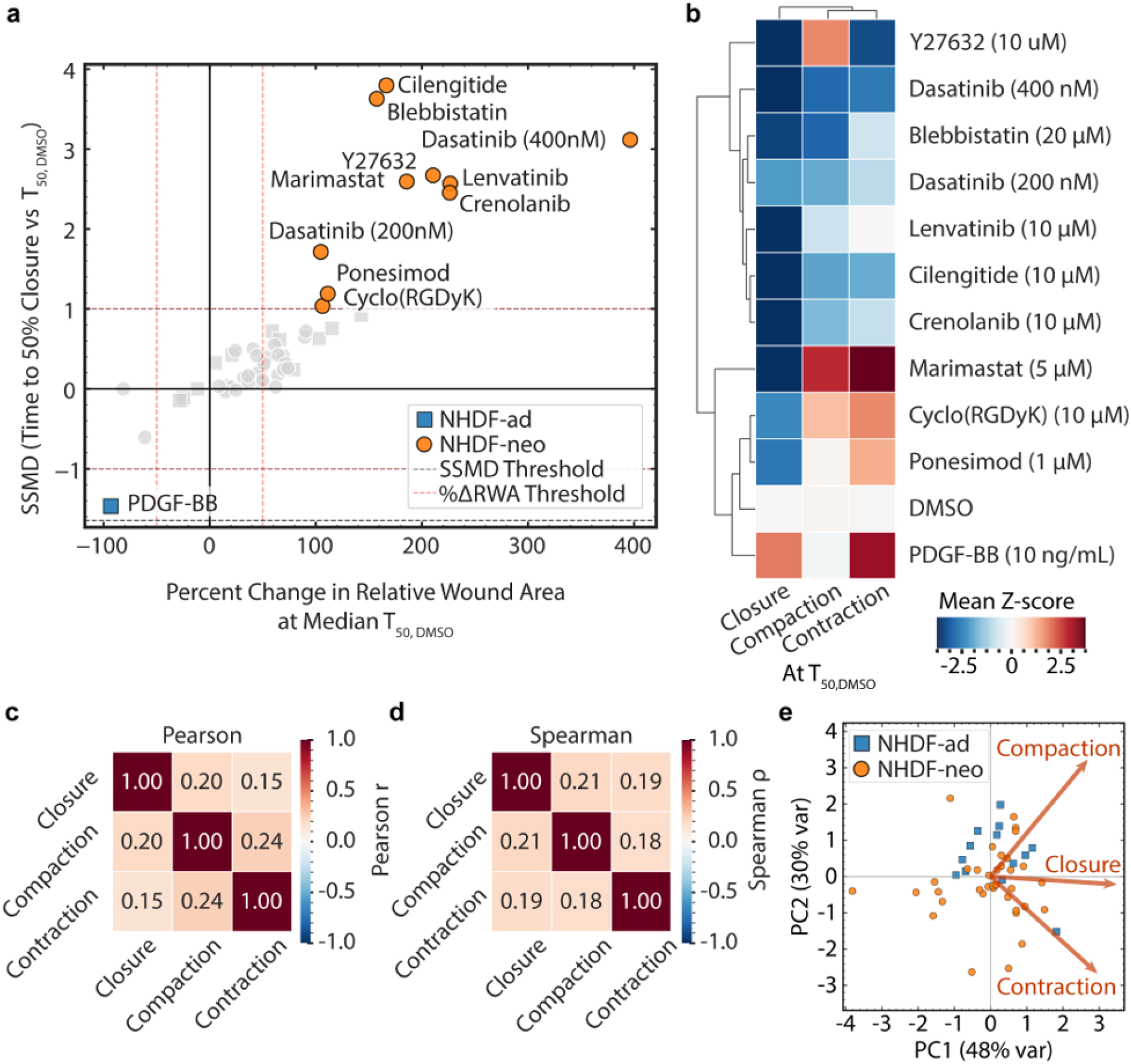
Multiparametric phenotyping of tissue remodeling. **(a)** Dual-flashlight plot with strictly standardized mean difference (SSMD) of time to 50% wound closure vs percent change in relative wound area at the time to 50% of the DMSO condition. Circles represent pharmacological compounds tested on NHDFneo cells and squares represent compounds tested on NHDFAd cells. Orange circles and blue squares represent hits that are above the designated thresholds (SSMD ±1 and %RWA ±50%). **(b)** Hierarchically clustered heatmap of DMSO-referenced z-scores across the three readouts: closure (1-relative wound area at T_50, DMSO)_, compaction (-% change in tissue footprint) at T_50, DMSO,_ and the change in pillar force at T_50, DMSO w_ith respect to pre-injury pillar force. Dendrogram representing the hierarchical clustering using Euclidean distance (Ward’s method) is displayed. Red/blue = increase/decrease relative to vehicle (mean z-score at T₅₀, DMSO. **(c)** Pearson linear correlation coefficients and **(d)** Spearman rank correlation coefficients for the three quantified metrics across all tested compounds and concentrations (n = 49). **(e)** PCA biplot of the condition z-score profiles (49 conditions). The direction and magnitude of the orange arrows represent the contribution of each metric to the dataset’s variance orange circles: NHDFneo and blue squares: NHDFAd cells.

Although several compounds produced comparable closure kinetics, inspection of tissue compaction and contractile force trajectories suggested that similar closure outcomes arose from distinct mechanical remodeling states. To systematically interrogate relationships between remodeling metrics, we compared closure kinetics, tissue compaction, and contractile force generation across all screened conditions. Hierarchical clustering of z-scored remodeling parameters, including relative gap area at T_50, DMSO (_gap closure), relative tissue footprint at T_50, DMSO (_tissue compaction), and change in contractile force at T_50, DMSO n_ormalized to pre-injury baseline, revealed distinct phenotypic response classes (Fig. 4b). Correlation analyses across screened conditions confirmed weak coupling between gap closure, compaction, and contractile force (maximum Pearson |r| ≤ 0.4) (Fig. 4c, d). Principal component analysis (PCA) further demonstrated that the variance across these phenotypic responses could not be captured by a single dominant axis; the first principal component (PC1) accounted for only 48% of total variance and the PC2 accounted for 30% of the variance (Fig. 4e). Evaluation of the PCA loadings revealed that gap closure and contractile force were weakly coupled to one another, while tissue compaction diverged along a distinct, partially opposed axis. These findings support the concept that gap closure, tissue compaction, and contractile force represent independent variables of mechanical tissue remodeling rather than intrinsically coupled outputs of repair. Together, these analyses indicate that remodeling outcomes cannot be reliably inferred from any single metric, highlighting the value of simultaneous multiparametric measurement.

### 3D microtissue closure aligns with clinical observations

To determine whether the mechanically coupled 3D microenvironment better predicts clinically observed therapeutic responses than conventional methods, we evaluated a panel of clinically documented compounds in both our microtissue platform and standard two-dimensional scratch assays (Fig. 5). The 3D microtissue platform detected inhibitory effects of clinically relevant compounds, including dasatinib^41^, nintedanib^41^, cisplatin^45^, and semaglutide^46^, all of which failed to produce measurable responses in the 2D assay (Fig. 5a-c). Both models correctly identified the pro-reparative activity of PDGF-BB^39^, whereas neither assay detected reported benefits of Metformin^47,48^.

**Fig. 5.**
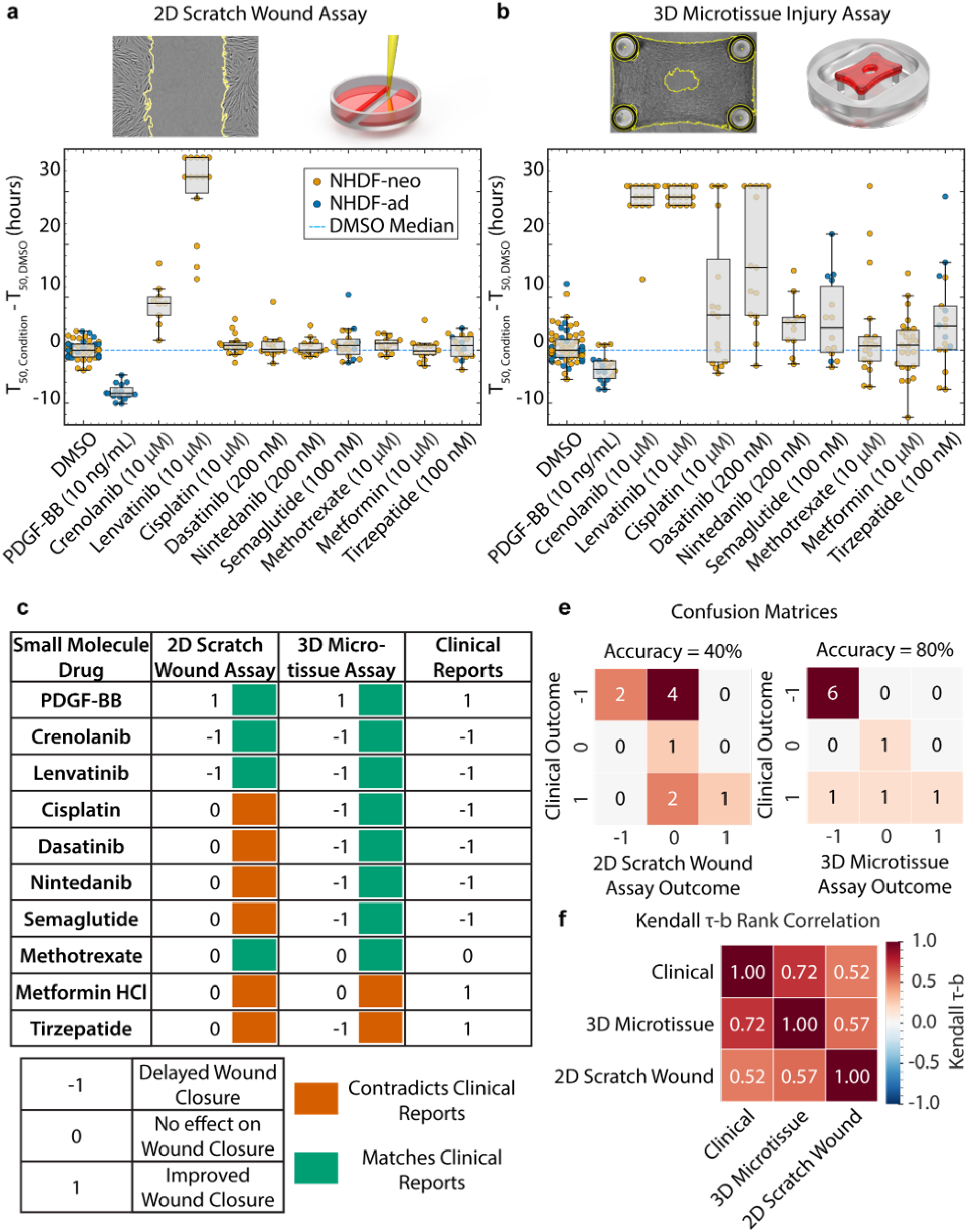
Benchmarking against planar scratch-wound assay. **(a)** 2D scratch wound assay with per-condition change in time-to-50%-closure relative to each plate’s median DMSO (T₅₀,condition − T₅₀,DMSO, h) **(b)** 3D microtissue repair assay with per-condition change in time-to-50%-closure relative to each plate’s median DMSO (T₅₀,condition − T₅₀,DMSO, h). Data from individual tissues are plotted as blue and orange dots representing NHDF-Neo and NHDF-Ad cells. (N=3) **(c)** Qualitative scoring of wound closure outcomes in clinical data, microtissue assays, and conventional scratch wound assays. Effects are categorized as delayed (–1), neutral (0), or accelerated (+1) wound closure. Compound effects matching clinical reports were annotated with a green square and effects contradicting clinical reports were annotated with red squares. **(d)** Classification of performance of the 2D Scratch Wound Assay and 3D Micro-tissue Assay (right) when evaluated against the ground truth established by clinical reports (n=10) as a confusion matrix. **(e)** Pairwise Kendall τ-b rank correlation coefficients between the 2D Scratch Wound Assay, 3D Micro-tissue Assay, and Clinical Reports (Ground Truth) for this set of small molecules (n=10).

Importantly, the 2D assay did not identify any clinically relevant responses not captured by the 3D platform (Fig. 5c, d). Kendall τ-b and Spearman’s rank correlation analysis quantified concordance between experimental outcomes and clinical observations (Fig. 5e, Supplementary Fig 4). Closure responses measured in the 3D microtissue model demonstrated a marginally statistically significant correlation with clinical findings (Kendall τ-b = 0.72, *p*=0.045, Spearman correlation of *ρ*=0.75*, p*=0.045), whereas the 2D scratch assay did not reach statistical significance (Kendall τ-b = 0.52, *p*=0.25, Spearman correlation of *ρ*=0.55*, p*=0.25). Together, these results demonstrate that closure dynamics measured in 3D microtissues provide improved functional alignment with clinically observed remodeling responses compared with conventional planar assays.

## III. DISCUSSION

Tissue repair is coordinated by dynamic interactions between cells, ECM, and mechanical forces, yet most scalable assays reduce this process to a single endpoint: gap closure. This simplification obscures how tissues remodel, transmit force, compact matrix, and recover structural continuity. To address this gap, we developed a 96-well microtissue platform that combines micropillar-based force measurement with tissue-scale imaging to quantify stromal remodeling in a format that is both mechanically informative and experimentally scalable. By integrating reusable 3D-printed modules, standard phase-contrast microscopy, and an open-source analysis pipeline, the platform provides access to assessing tissue remodeling processes that are difficult to capture in conventional 2D assays and impractical to measure at scale in existing 3D systems.^49–51^

A key component of this workflow is WoundCompute, an open-source computational framework for automated analysis of microtissue remodeling across large experimental datasets. Existing image-analysis tools often require proprietary software, or have limited batch-processing automation, insufficient documentation, and inaccessible source code, creating barriers to reproducibility and adoption.^52–60^ Moreover, supervised learning approaches introduce an additional constraint i.e. when manual annotations are treated as ground truth, model accuracy is inherently bounded by the consistency of human labeling. As shown in Appendix S1, identical instructions can yield substantial annotator-to-annotator variation, therefore limiting reproducibility across researchers and laboratories.^61–63^ WoundCompute avoids this dependency by using deterministic image-analysis routines, enabling reproducible, large-scale quantification without manual intervention.

Multiparametric phenotyping revealed that closure, compaction, and contractile forces are related but separable features of tissue repair. Hierarchical clustering of pharmacological perturbations identified distinct remodeling signatures associated with contractile motor activity, adhesive coupling, and proteolytic matrix remodeling. Inhibition of the non-muscle myosin II motor with blebbistatin, which blocks ATPase activity downstream of both the Rho-kinase (ROCK) and myosin light chain kinase (MLCK) pathways, simultaneously impaired closure, contractile force generation, and tissue compaction.^64,25,29,42^ This phenotype is consistent with disruption of the common actomyosin machinery that drives both force generation and fibrillar matrix densification. Dasatinib produced a similar reduction in contractile tissue force, compaction and closure, but its interpretation is less straightforward. As a broad-spectrum tyrosine kinase inhibitor targeting Src-family kinases, Abl, PDGFR,^65,66^ it also suppresses ECM production, including collagen synthesis^67^, making it difficult to attribute this phenotype to a single molecular target.

In contrast, selective ROCK inhibition with Y-27632 impaired closure with reduced tissue contractile force but increased tissue compaction. This phenotype suggests that relieving global stress-fiber tension while preserving MLCK-dependent myosin activity allows local collagen densification and cell packing to continue despite diminished tissue-scale forces. Alternatively, increased compaction in the presence of impaired closure may reflect altered focal adhesion turnover and matrix remodeling that are not captured by measurement of contractile force alone.^68,69^ Integrin inhibition with cilengitide impaired closure with minimal effects on contractility or compaction, consistent with an adhesion-mediated failure to transmit mechanical work effectively to the gap interface.^43,70^ These results indicate that tissue contractility and compaction cannot be interpreted as a proxy for closure. Rather, efficient tissue repair depends on the coordinated regulation of force production, cell-matrix adhesion, and matrix reorganization, rather than on maximizing any single mechanical output.

This distinction is important for interpreting therapeutic responses. Closure measured in microtissues showed stronger agreement with reported clinical outcomes than conventional planar scratch assays, likely because 3D stromal repair depends on cell–matrix coupling, matrix remodeling, and tissue-scale force redistribution.^71^ Planar assays primarily capture cell migration across rigid substrates and lack the dynamic matrix remodeling and tissue-scale force transmission that regulate stromal repair in vivo.^71,72^ Consequently, pharmacological compounds that modulate matrix remodeling, or mechanical coupling may show limited to no effect in 2D systems.^71^ The microtissue platform also captures an early phase of gap expansion following injury. This transient widening likely reflects release of tissue prestress, elastic recoil and clearance of the damaged matrix after disruption of mechanical continuity across the wound.^27^ Quantifying this phase extends the assay beyond closure-only readouts and increases sensitivity to both repair-promoting and repair-inhibiting responses.

A major limitation of the current model is its focus on fibroblast-driven stromal remodeling. Fibroblasts are central regulators of matrix contraction, deposition, and repair mechanics, but in vivo wound repair also depends on epithelial, immune, endothelial, and perivascular cell populations that regulate inflammation, angiogenesis and barrier restoration.^73–75^ As a result, the remodeling signatures identified here should be interpreted as stromal mechanical phenotypes rather than complete wound-healing programs. Incorporating these additional cell types into the platform will be necessary to determine how multicellular signaling modifies force generation, matrix remodeling, and closure dynamics.

The platform is well positioned for future studies that combine genetic perturbation with tunable matrix composition to systematically define how specific ECM components regulate productive repair and pathological remodeling. Coupling mechanical phenotyping with molecular profiling would further enable multi-scale mapping of the regulatory networks governing tissue remodeling.^76^ By separately quantifying tissue closure, matrix compaction, and contractile force generation, this platform establishes a generalizable experimental framework for phenotyping functional tissue remodeling processes across diverse biological contexts.

## IV. METHODS

### Mold Fabrication and Assembly

To achieve both the high-fidelity resolution required for microscale pillar force-sensor templating and the modularity necessary for scalable handling, we developed a hybrid fabrication strategy combining two-photon polymerization (2PP) and stereolithography (SLA) (Supplementary Fig. 1A). We utilized 2PP to fabricate a master mold containing inverse cavities for a four-pillar array (1 mm x 0.6 mm) situated within a 1.12 × 1.5 mm microwell (Supplementary Methods). Each pillar features a 150 µm diameter cylindrical base, a 200 µm diameter spherical cap, and a total height of 450 µm (Fig. 1B). To scale production without degrading the original master, we established a replica molding pipeline and fabricated a 4x5 configuration epoxy resin (SmoothCast 322, Smooth-On) replica (Supplementary Methods). Multiple copies of the master mold were then fabricated in polydimethylsiloxane (PDMS, 1:10 Sylgard 184, Dow) to be used as our working PDMS stamps. To adapt these stamps for high-throughput formats, individual PDMS stamps were secured into custom SLA-printed holders using a twist-lock cap mechanism. Assembling these discrete units onto SLA-printed racks yielded an 8×12 mold array with standard 9 mm spacing. This assembly is then coated with trichloro(1H,1H,2H,2H-perfluorooctyl) silane for further downstream use.^77,78^ Because this architecture is highly modular, individual stamps can be replaced and silanized again without sacrificing the entire array, enabling the platform to be reused for more than 30 cycles with minimal maintenance.

### Plate fabrication

To mitigate edge effects driven by uneven evaporation, we utilized 96-well plates featuring a PBS-fillable perimeter moat (Nunc Edge 2.0 96-well plate, Thermo Scientific, Cat No. 267578).^79^ Within each well, PDMS devices were fabricated by vacuum-casting PDMS onto the assembled 8×12 mold array, inserting the molds into the plate, and curing at 60 °C for at least 6 hours.^77,78^ Once de-molded, the plate was sonicated with 70% ethanol in each well to clean the fabricated devices, then air dried and stored for later use.

### Surface treatment optimization

To direct microtissue attachment to the pillar caps while preventing spurious cell adhesion and matrix assembly in the microwells, we systematically evaluated six surface treatment strategies (Table 1). These strategies explored varying sequential combinations of a polydopamine (PDA) adhesive layer (1 mg/mL in 1X TBS, pH 8.5), poly(L-lysine)-graft-poly(ethylene glycol) (PLL-g- PEG) (1 mg/mL in 10 mM HEPES Buffer, pH 7.4), and Pluronic F-127 (in water). The optimized protocol (Strategy 6) consisting of global PDA coating, localized PLL-g-PEG passivation below the caps, overnight aqueous incubation at 37 °C to minimize bubble formation during tissue culture, and a final 2% Pluronic F-127 backfill yielded the most robust tissues and was employed for all subsequent experiments. Within this protocol, the stable PDA/PLL-g-PEG coating prevents fouling, while the F-127 backfill covers regions left exposed by the steric hindrance of large PEG molecules. These functionalized wells were sterilized by a 70% ethanol rinse followed by 15 minutes of ultraviolet irradiation prior to cell seeding.

| SN | Strategy Name | Layer 1 | Wash<br>(H<sub>2</sub>O) | Layer 2 | Wash<br>(H<sub>2</sub>O) | Equilibration | Layer 3 | Wash<br>(H<sub>2</sub>O) |
| --- | --- | --- | --- | --- | --- | --- | --- | --- |
| 1 | 2% F-127 | 2% Pluronic F-127,<br>50 µL, 10 mins, 25° C | 1X |  |  | Sterile water, 37° C,<br>overnight |  |  |
| 2 | 2% F-127+PDA | 2% Pluronic F-127,<br>50 µL, 30 mins, 25° C | 1X | 1 mg/mL PDA,<br>50 µL, 30 mins, 25° C | 6X | Sterile water, 37° C,<br>overnight |  |  |
| 3 | PDA+2% F-127 | 1 mg/mL PDA,<br>50 µL, 30 mins, 25° C | 6X | 2% Pluronic F-127,<br>50 µL, 30 mins, 25° C | 1X | Sterile water, 37° C,<br>overnight |  |  |
| 4 | 40% F-127+PDA | 40% F-127,<br>0.65 µL, 30 mins, 4° C | No | First incubate plate, 37°<br>C, 10 mins to gel F-127;<br>add 1 mg/mL PDA,<br>50 µL, 30 mins, 37° C,<br>layered over F-127 gel | 6X<br>(cold) | Sterile water, 37° C,<br>overnight |  |  |
| 5 | PDA+PLL-g-PEG | 1 mg/mL PDA,<br>50 µL, 30 mins, 25° C | 6X | 1 mg/mL PLL-g-PEG,<br>0.65 µL, 1 hour, 25° C | 3X | Sterile water, 37° C,<br>overnight |  |  |
| 6 | PDA+PLL-g-PEG<br>+2% F-127 | 1 mg/mL PDA,<br>50 µL, 30 mins, 25° C | 6X | 1 mg/mL PLL-g-PEG,<br>0.65 µL, 1 hour, 25° C | 3X | Sterile water, 37° C,<br>overnight | 2% Pluronic F-127,<br>50 µL, 10 mins, 25°<br>C | 1X |

### Cell culture

Primary normal human dermal fibroblasts from neonatal foreskin (NHDF-Neo, Lonza) or from adult skin (NHDF-Ad, Lonza) were expanded in fibroblast growth medium 2 (FGM2, Lonza) at 37° C, 5% CO_2 a_ccording to manufacturer instructions. Passages 2-6 were used in our experiments.

### Microtissue formation and tissue repair model

To generate 3D microtissues, cells were suspended in neutralized liquid rat tail type I collagen (2.2 mg/mL, Corning) and seeded into each device in 0.9 µL volumes. To prevent premature matrix polymerization during high-throughput array loading, plates were maintained on ice throughout the seeding process. Seeding densities were titrated between 2,000 and 12,000 cells per well for optimization and a standard density of 4,000 cells per well was established for all subsequent experiments. Following loading, devices were inverted and incubated at 25 °C for 15 minutes, followed by 37 °C for 15 minutes to ensure uniform collagen polymerization. Finally, each well was supplemented with FGM2 containing 50 µg/mL L-ascorbic acid-2-phosphate, and cells were allowed to drive tissue assembly overnight at 37 °C.

To initiate tissue repair and remodeling, laser ablation was used to incite a full-thickness injury as previously described^27^. Briefly, a single, precise injury was ablated in the center of each 3D stromal tissue using a q-switched nanosecond-pulsed Nd:YAG laser (Minilite I, Continuum) emitting at 1064 nm wavelength with 1 mJ of power.

### Time-lapse microscopy

Phase contrast images of microtissues were first captured at 10X before ablation on an automated Nikon Ti Eclipse microscope fitted with a Lumenera Infinity 8-2M camera and an environmental chamber (37 °C, 5% CO_2)_. Microtissues were then ablated as described. Phase contrast, time-lapse images of ablated microtissues were then taken every 30 min for at least 24 hours.

### WoundCompute processing, manual processing and measurements

WoundCompute is software designed to quantify wound area, tissue footprint, and pillar positions automatically at every imaging timepoint; full details of the WoundCompute segmentation and tracking pipeline are provided in Appendix S1. Briefly, individual micropillars were segmented from phase contrast images using Otsu thresholding and morphological operations (i.e., closing, dilation, erosion, and opening), with pillar cross-sections identified via ellipse fitting and clustering. Pillar positions were tracked across frames using masked template matching with normalized cross-correlation. Wound and tissue segmentation were performed through a separate pipeline comprising Gabor filtering, median and Gaussian smoothing, and Otsu thresholding, where the combined pillar mask defined an admissible region within which the wound was identified as the largest connected component. The tissue mask was obtained by inverting the thresholded image and retaining the largest connected component not touching the image boundary. The pipeline is implemented as a modular, test-driven framework in which each processing step is independently validated through unit tests. Pillar tracking accuracy was validated by applying synthetic displacements of known magnitude, and the recovered synthetic displacements were assessed against the known ground truth. Segmentation was evaluated by comparing WoundCompute extracted wound and tissue area against manual annotations on a subset of images, where manual segmentation served as a reference check rather than a ground-truth benchmark. WoundCompute runs on all three major operating systems (Microsoft Windows, MacOS, and Linux) and is openly available under the MIT License.

For all manual validation, wound area and tissue footprint were measured every 2 hours in ImageJ using the polygon selection tool. Pillar coordinates were tracked using the centroid of the ellipse tool. Details on validation and variability between manual annotators are provided in Appendix S1

To yield the relative wound area (*RWA*), the wound area was normalized to the initial measurement recorded at the first timepoint post-injury. The tissue footprint was normalized to the initial tissue footprint prior to injury to calculate the percent change in relative tissue footprint (*RTF%*).

The change in average pillar force (*RPF*) was then calculated as previously described.^25,28,29^ Briefly, the sum of pillar displacements (*d*) at each time point were calculated by tracking the shift in pillar centroids relative to their pre-injury positions (Appendix S1). We calculated the change in force using Hooke’s law (*FF* ≈ *kk* × *dd*) using an effective spring constant (*k*) of 2 μN μm^−1^ (Supplementary Fig. 6) for a typical Young’s modulus of 1.7 MPa for 1:10 PDMS cured at 60 °C.^80^

### High-throughput screen

Microtissues were seeded in multi-well plates and allowed to self-assemble as previously described. Small molecule targets or vehicle control (0.1% DMSO) were assigned to randomized wells and added to the culture media immediately prior to injury. A comprehensive list of the evaluated compounds and their corresponding working concentrations selected to be 5-10X of the IC_50 i_s provided in Supplementary Table 1. Following treatment, time-lapse microscopy, laser ablation, and automated quantification via WoundCompute were executed across all arrays utilizing the established protocols.

### Scratch Wound Assay

The scratch wound assay was performed as described in ^17,81^. Briefly, human dermal fibroblasts (NHDF-Neo or NHDF-Ad) were seeded in a 96 well plate at a density of 20,000 cells/well in FGM2 media supplemented with 50 ug/mL L-ascorbic acid-2-phosphate. Cells were grown to 95% confluency at 37°C and 5% CO_2._ A linear scratch was created in each monolayer using an Agilent BioTek AutoScratch device. Wells were washed twice with PBS to remove dislodged cells and fresh media containing either vehicle (0.1% DMSO) or one of the small molecules with reported clinical outcomes: Crenolanib (10 µM), Cisplatin (10 µM), Dasatinib (200 nM), Lenvatinib (10 µM), Metformin HCl (10 µM), Methotrexate (10 µM), Nintedanib(200 nM), PDGF-BB (10 ng/mL), Semaglutide (100 nM), Tirzepatide (100 nM).

Images of the wound were captured immediately following injury and every 30 minutes thereafter for 48 hours using an Agilent BioTek Cytation 10 microscope (10X objective). The wound area was quantified at each time point using Gen5 Image Prime software (Agilent BioTek), and wound closure was normalized to the initial wound area.

#### Statistics

All standard statistical analyses were performed in Python 3 (SciPy and scikit-learn). Prior to analysis, data distribution normality was assessed using the Shapiro-Wilk test, and variance equality was confirmed via the F-test (for two groups) or the Brown-Forsythe test (for multiple groups). Outliers were identified using the Grubbs’ test (α = 0.05) and excluded before normality was re-evaluated. For comparisons between two independent groups, normally distributed data were analyzed using an unpaired, two-tailed Student’s *t*-test (equal variance) or Welch’s *t*-test (unequal variance). Non-normally distributed data were evaluated using the Mann-Whitney *U* test. Statistical significance was defined as *P* < 0.05.

For the high-throughput and scratch-wound screens, individual wells were filtered before analysis using an interquartile-range outlier rule (values below Q1 − 1.5 × IQR or above Q3 + 1.5 × IQR at the baseline timepoint) together with WoundCompute tissue-breakage and manual quality-control evaluations.

To quantify the effect of each compound in the high-throughput screen, the strictly standardized mean difference (SSMD)^38,82^ of the time to 50% relative wound area (*T_50_*_)_ computed for each cell type – condition pair against its same-plate DMSO control:

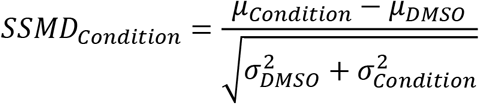

*Where μ is the mean and σ^2^ is the variance of T_50_*

Hits were identified on a dual-flashlight plot^83^, where we plotted the SSMD (y-axis) against the percentage change in relative wound area evaluated at *T_50_* _o_f DMSO-treated control tissues (*T_50, DMSO_*_)_ (x-axis). Compounds with SSMD > ±1 and percentage change in relative wound area > ±50% at *T_50, DMSO_* _w_ere classed as moderate to strong hits. Here, a positive SSMD indicates a longer T50 (delayed wound closure) relative to the same-plate DMSO control, whereas a negative SSMD indicates accelerated closure.

For comparative profiling across multiple parameters, standard scores (z-scores) for closure, compaction and contraction were respectively calculated as the change in relative wound area (1-*RWA*), the negative percent change in tissue footprint relative to the pre-injury baseline (-*RTF%*), and the change in pillar force relative to pre-injury pillar force (*PF*), all evaluated at *T_50, DMSO_*. Each metric was standardized against the same-plate DMSO control distribution. The z-scores for these metrics correspond to the Closure, Compaction, and Contraction axes and were calculated as:

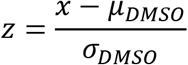

*Where x is the value of the metric measured from a well μ_DMSO_ is the mean and σ_DMSO_ is the standard deviation of the DMOS metric*

Condition-level z-scores (mean over wells) were clustered hierarchically (Ward’s linkage, Euclidean distance; SciPy^84^) to generate a phenotypic heatmap. Principal-component analysis was performed on the condition-level z-score matrix (closure, compaction, contraction) using scikit-learn (mean-centered and unit-variance scaled). Pearson and Spearman correlations among the condition-mean z-scores were computed, with off-diagonal cells marked at p < 0.05.

For the by-condition analyses of 3D microtissue closure and scratch wound closure, each well’s relative wound-area trajectory was reduced to the time to 50% closure (T50) and expressed as ΔT50 relative to the same-plate median DMSO T50; wells that did not reach 50% closure within the imaging window were assigned that plate’s maximum imaging time before normalization.

Finally, to benchmark clinical concordance, outcomes from both published clinical reports and the in vitro data (based on the measured *T_50_*_)_ were discretized into a scoring system: For each assay, a compound’s rating was taken from the cell-type/condition with the strongest effect (largest |SSMD| of T50): −1 (delayed) or +1 (accelerated) when |SSMD| ≥ 0.7, and 0 (no effect) otherwise, with the SSMD oriented so that closure faster than the same-plate DMSO control is positive (matching the clinical +1 = accelerated convention). (Supplementary Table 3 and Supplementary Table 4). Agreement between the clinical and assay ratings (n = 10 conditions) was quantified with Spearman’s rank correlation (*ρ*) and Kendall’s *τ-b.* The significance of the Spearman *ρ* and Kendall *τ-b* coefficients was assessed with a two-sided permutation test (9999 random permutations of the paired ratings, with a fixed random seed for reproducibility). Agreement was additionally visualized as 3×3 confusion matrices of each in-vitro assay (engineered microtissue and 2D scratch-wound) against the clinical calls.

## Supporting information

Appendix-S1: WoundCompute

Appendix-S2: Published Dataset Structure

## ACKNOWLEDGEMENTS

We would like to thank Dr. Christopher S. Chen for his insightful discussions; Dr. Christos Michas for insightful discussions, help fabricating prototypes and training; Dr. Megan Griebel for her insights and training; Soham Kulkarni for his technical assistance; and Laurie Kelleher and Amy B Michaels for administrative and technical support. Additionally, we would like to thank the Biological Design Center and Photonics Center at Boston University for additional technical support and instrumentation access.

## AUTHOR DECLARATIONS

### Ethics Approval

Ethics approval is not required.

### Author Contributions

ASV, QBN, EL and JE conceived this study. ASV and QBN designed and analyzed the experiments. ASV, QBN, ED, MÇK, EW, and VS, performed the experiments. ASV, QBN, EL and JE wrote this manuscript with feedback from all authors. JE, EL and WW supervised and funded this study.

## DATA AVAILABILITY

The data that support the findings of this study are available on BioImage Archive (10.6019/S-BIAD2719).

## CODE AVAILABILITY

WoundCompute source code is available at https://github.com/elejeune11/woundcompute, and a graphical interface for noncomputational users is available at https://github.com/quan4444/woundcomputeGUI. Further documentation for both is provided in the respective README files and in Appendix S1 (WoundCompute Software). WoundCompute is archived on Zenodo (<u>10.5281/zenodo.17127155</u>).

