## Supplementary Materials and Methods for "A high-throughput, 3D microtissue platform for multiparametric analysis of tissue remodeling"

### SUPPLEMENTARY FIGURES

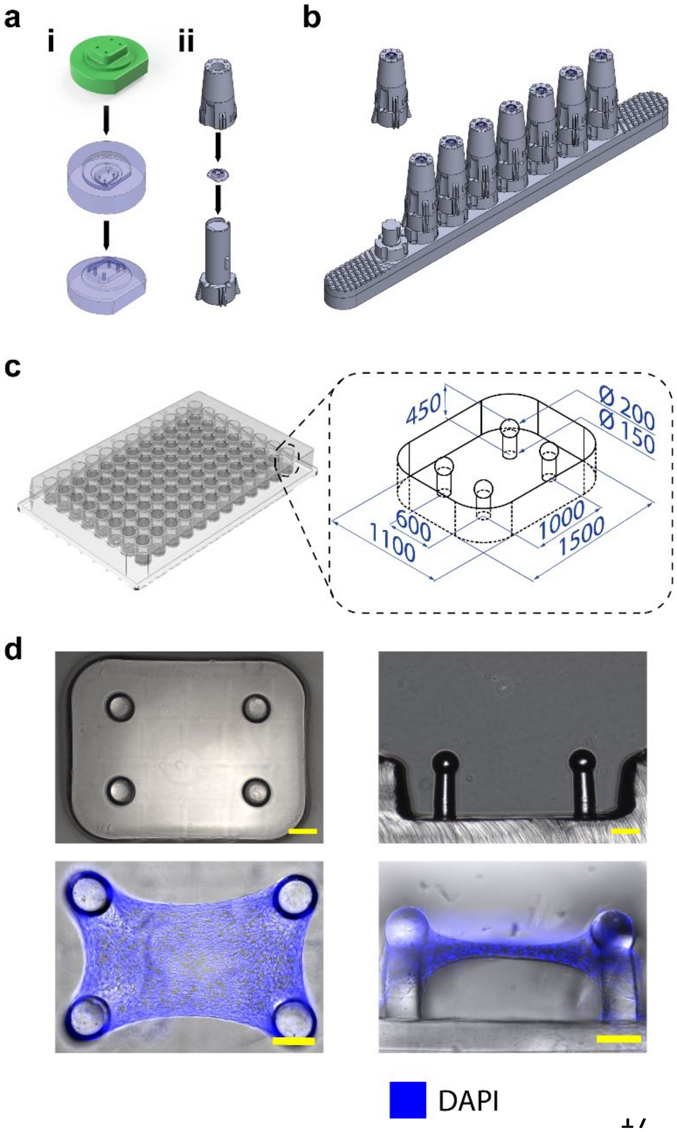


**Supplementary Fig. 1. Assembly of 3D-printed mold. (a)** Representative images of the assembly of the molds and mold-holder system. i) 2 photon-polymerization master molds of the pillar devices copied to PDMS to create PDMS stamps. ii) Insertion of PDMS stamp in 3D printed mold-holder and secured with a twist-lock mechanism. **(b)** Eight mold holders assembled onto aligning rack with twist-lock mechanisms. **(c)** Schematic diagram of the micropillar array in a well with dimensions (µm). **(d)** Phase contrast images of the top and cross-sectional view of the devices and tissues (before and after injury) are shown. DAPI channel overlaid. Scale bar: 200 µm.

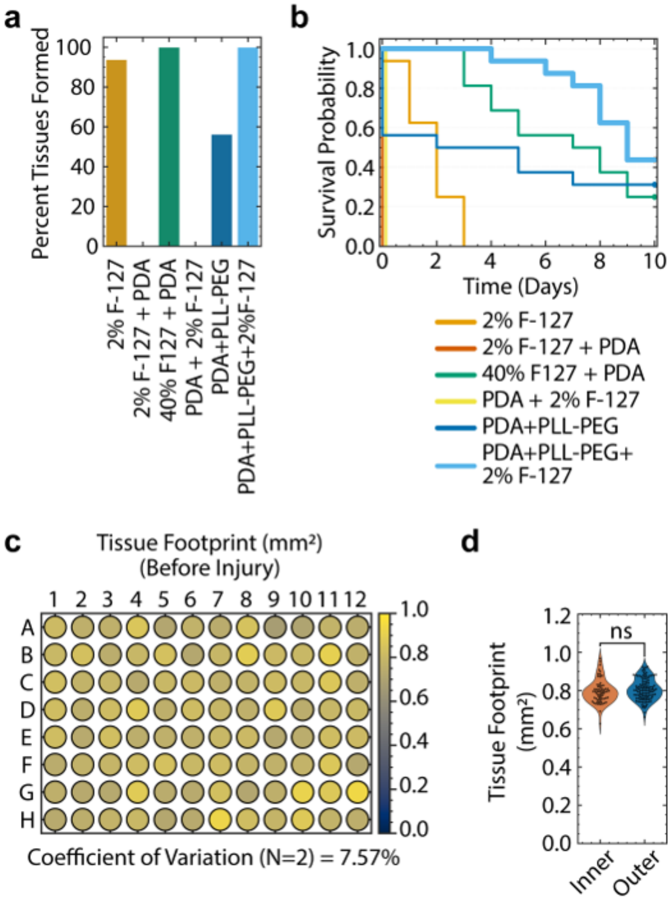


**Supplementary Fig. 2. Parameter optimization for robust ECM-remodeling assays. (a)** Percent tissues that successfully self-assembled on each type of surface treatment. (N=1, n=12) **(b)** Kaplan-Meier plot representing tissue survival on all surface treatments to alter the cell-adhesive properties of native PDMS. Tissue survival is defined as the number of days a tissue remained attached to all 4 pillars. (N=1, n=12). F-127: Pluronic F-127, PDA: Polydopamine-HCl, PLL-PEG: poly-l-lysine-g-polyethylene glycol block copolymer. **(c)** Heatmap representing average tissue footprint in each well at $T_0$ (N=2). Tissue footprint is represented from 0 to 1 $mm^2$ from white to green **(d)** Tissue area at $T_0$ grouped by location (Outer wells: all wells located in Row A, H and Columns 1, 12; Inner wells: wells 2B to 11G). ns: not significant, unpaired t-test.

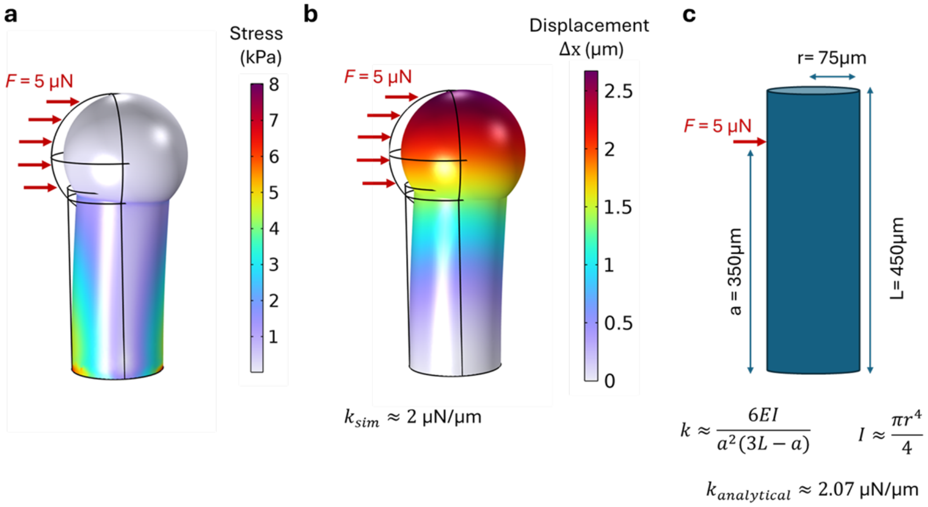


**Supplementary Fig 3. Calculation and simulation of effective spring constant**

Finite element modeling of the devices implemented using COMSOL Multiphysics with (a) stress (b) displacement due to the force applied at the spherical cap of the pillars. (c) Analytical approximation of the effective spring constant.

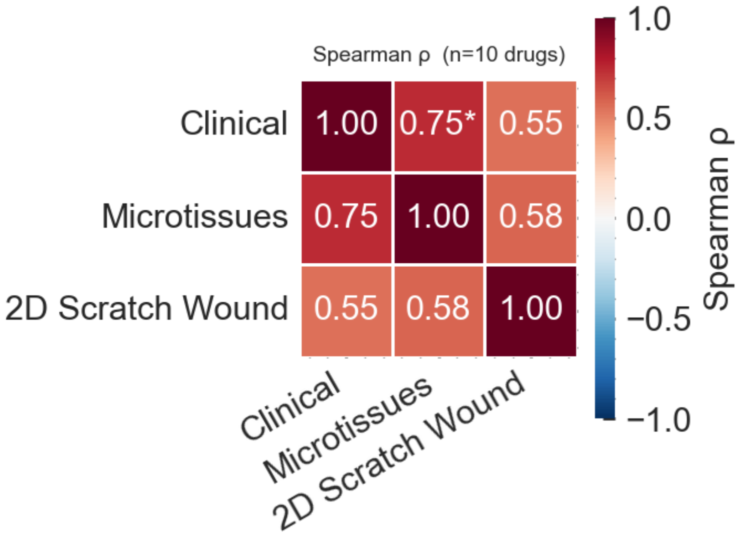


**Supplementary Fig 4 Calculation of the Spearman ranked correlation of clinical outcomes, 2D scratch wound assay and 3D microtissue assay.** Coefficients between the 2D Scratch Wound Assay, 3D Micro-tissue Assay, and Clinical Reports (Ground Truth) for this set of small molecules (n=10)

#### SUPPLEMENTARY TABLES

##### Supplementary Table 1. Pharmacological compounds and concentrations tested

| S.N. | Compound Name | Concentration | Source | Catalog Number |
|---|---|---|---|---|
| **1** | AICAR | 20 µM | APEXBio DiscoveryProbe™ FDA-approved Drug Library (L1201) | A8184 |
| **2** | Blebbistatin | 20 µM | Cayman Chemical | 13013 |
| **3** | Cilengitide | 10 µM | APEXBio DiscoveryProbe™ FDA-approved Drug Library (L1201) | A8660 |
| **4** | Cisplatin | 10 µM | APEXBio DiscoveryProbe™ FDA-approved Drug Library (L1201) | A8321 |
| **5** | Crenolanib (CP-868596) | 10 µM | APEXBio DiscoveryProbe™ FDA-approved Drug Library (L1201) | A8307 |
| **6** | Cyclo(RGDyK) | 10 µM | APEXBio DiscoveryProbe™ FDA-approved Drug Library (L1201) | B6015 |
| **7** | Dasatinib | 200, 400 nM | APEXBio DiscoveryProbe™ FDA-approved Drug Library (L1201) | A3017 |
| **8** | Dinaciclib (SCH727965) | 5 µM | APEXBio DiscoveryProbe™ FDA-approved Drug Library (L1201) | A8412 |
| **9** | EMD-1214063 | 100 nM | APEXBio DiscoveryProbe™ FDA-approved Drug Library (L1201) | A3388 |
| **10** | Exendin-4 | 1, 10, 100 nM | MedChemExpress | HY-13443 |
| **11** | Fingolimod (FTY720) | 100 nM | APEXBio DiscoveryProbe™ FDA-approved Drug Library (L1201) | A8548 |
| **12** | Lenvatinib | 10 µM | Lenvatinib (E7080) | A2174 |
| **13** | Liraglutide | 1, 10, 100 nM | Cayman Chemicals | 24727 |
| **14** | Marimastat | 5 µM | APEXBio DiscoveryProbe™ FDA-approved Drug Library (L1201) | A4049 |
| **15** | Metformin HCl | 10 µM | APEXBio DiscoveryProbe™ FDA-approved Drug Library (L1201) | B1970 |
| **16** | Methotrexate | 10 µM | APEXBio DiscoveryProbe™ FDA-approved Drug Library (L1201) | A4347 |
| **17** | Nintedanib (BIBF 1120) | 200 nM | APEXBio DiscoveryProbe™ FDA-approved Drug Library (L1201) | A8252 |
| **18** | PDGF-BB | 10, 100 ng/uL | R & D Systems | 220-BB-010 |
| **19** | Ponesimod | 1 µM | Cayman Chemical | 22053 |
| **20** | Quercetin Dihydrate | 10 µM | APEXBio DiscoveryProbe™ FDA-approved Drug Library (L1201) | A2552 |
| **21** | Quercetin Dihydrate + Dasatinib | 10 µM + 200 nM | APEXBio DiscoveryProbe™ FDA-approved Drug Library (L1201) | A2552 + A3017 |
| **22** | Ruxolitinib (INCB018424) | 1 µM | APEXBio DiscoveryProbe™ FDA-approved Drug Library (L1201) | A3012 |

| | | | | |
|---|---|---|---|---|
| **23** | SB431542 | 10 µM | Selleck Chemicals | S1067 |
| **24** | Semaglutide | 1, 10, 100 nM | Cayman Chemicals | 40231 |
| **25** | Sunitinib | 20 µM | APEXBio DiscoveryProbe™ FDA-approved Drug Library (L1201) | B1045 |
| **26** | Tirzepatide | 1, 10, 100 nM | Cayman Chemicals | 39748 |
| **27** | Y27632 | 10 µM | Cayman Chemical | 10005583 |

**Clinical evidence of drug efficacy**

Scoring key: **-1** = delayed wound closure; **0** = no effect; **+1** = improved wound closure

**Supplementary Table 2**. **Clinical Reports and Scoring of Small Molecules**

| **Drug** | **Score** | **Clinical Indication** | **Key Outcome / Evidence** |
|---|---|---|---|
| PDGF-BB (becaplermin) | +1 | Chronic neuropathic diabetic ulcers of the lower extremities; cutaneous wound repair | 50% vs 35% complete DFU closure at 20 wk, p=0.007. [1]<br>Pooled 4 RCTs (n=922): 39% higher closure probability, p=0.007. [2] |
| Crenolanib | -1 | Newly diagnosed FLT3-mutant AML | Generally not administered during major surgery[3] and mechanism based[4] |
| Lenvatinib | -1 | Thyroid Cancer (DTC, RAI-R DTC), Renal Cell Carcinoma (RCC), Hepatocellular Carcinoma (HCC), Endometrial Carcinoma (EC) | FDA label mandates hold ≥1 wk pre-elective surgery; resume ≥2 wk post-op after adequate healing (Lenvima USPI, NDA 206947).<br>Fistula/GI perforation 2% across 799 patients in pooled safety.<br>14.7% fistula/perforation in RR-DTC cohort. [5] |

| | | | |
|---|---|---|---|
| Cisplatin | -1 | Ovarian cancer, Sarcoma | Impairs (primarily during proliferative stage). Recommended hold 1 week prior to surgery, restart only after wound healing achieved. [6] |
| Dasatinib | -1 | Chronic Myeloid Leukemia (CML), Acute Lymphoblastic Leukemia (ALL) | Impairs (post-TKA dehiscence case). [7] |
| Nintedanib | -1 | Idiopathic Pulmonary Fibrosis (IPF), Systemic Sclerosis-associated ILD, Advanced malignancies | FDA Ofev label: may impair wound healing; hold for surgery.<br>Australian lung transplant cohort: bronchial anastomotic dehiscence 7.5% vs 2.2% (p=0.08, nonsignificant). [8] |
| Semaglutide | -1 | Type 2 Diabetes, Obesity (weight loss) | Impairs (non-diabetic): Non-diabetic post-bariatric body contouring: dehiscence 5.19% vs 2.78%, delayed healing 2.58% vs 1.21%, p<0.0001.[9]<br>Diabetic surgical cohort, 35,020 propensity-matched procedures: dehiscence relative risk 0.711 (95% CI 0.577-0.877), p=0.001. Authors explicitly note results are not |

| | | | |
|---|---|---|---|
| | | | generalizable to non-diabetic populations. [10]<br>DFU TriNetX cohort also showed lower dehiscence 0.26% vs 0.56%. [11]<br>Context-dependent. -1 only for non-diabetic post-bariatric body contouring |
| Methotrexate | 0 | Inflammatory Arthritis (RA, AS, PsA, JIA), SLE, Sarcoma, Carcinoma | Randomized Controlled Trial in Rheumatoid Arthritis orthopedic surgery: continued MTX 2% vs stopped 15% complication rate, p<0.003. [12]<br>ACR/AAHKS 2022 guideline: continue Methotrexate through elective Total Hip Arthroplasty/ Total Knee Arthroplasty. [13] |
| Metformin | +1 | Type 2 Diabetes, Traumatic Ulcers, Hypertrophic/Keloid Scars, Periodontitis | Benefit likely glycemic and anti-inflammatory.[14] |
| Tirzepatide | -1 | Type 2 Diabetes, Weight management | Impairs, (non-diabetic): TriNetX cohort excluding diabetics, smokers, PVD, and metabolic/endocrine disease. Pooled GLP-1 agonists (including tirzepatide) vs none, propensity-matched 30-day wound complications: panniculectomy 4.7% vs |

| | | | |
|---|---|---|---|
| | | | 2.7% (p=0.05); abdominoplasty 9.8% vs 3.6% (p=0.001); breast reduction 2.6% vs 1.3% (p=0.035). [15] |

**Supplementary Table 3**. **Clinical Reports and Scoring of Small Molecules**

The scoring key: **-1** = delayed wound closure; **0** = no effect; **+1** = improved wound closure was applied based on SSMD effect sizes ≥ 0.7.

| **Drug** | **Clinical Ordinal Score** | **2D SSMD** | **2D Ordinal Score** | **3D SSMD** | **3D Ordinal Score** |
|---|---|---|---|---|---|
| Dasatinib | -1 | -0.39 | 0 | -3.12 | -1 |
| Lenvatinib | -1 | -4.4 | -1 | -2.57 | -1 |
| Nintedanib | -1 | -0.21 | 0 | -0.75 | -1 |
| Crenolanib | -1 | -1.52 | -1 | -2.46 | -1 |
| Metformin HCl | 1 | -0.01 | 0 | -0.23 | 0 |
| Cisplatin | -1 | -0.31 | 0 | -0.78 | -1 |
| Methotrexate | 0 | -0.3 | 0 | -0.38 | 0 |
| PDGF-BB | 1 | 3.58 | 1 | 1.47 | 1 |
| Semaglutide | -1 | -0.63 | 0 | -0.92 | -1 |
| Tirzepatide | 1 | -0.69 | 0 | -0.72 | -1 |

#### SUPPLEMENTARY METHODS

##### Master Mold Fabrication

To achieve both the high-fidelity resolution required for microscale pillar force-sensor templating and the modularity necessary for scalable handling, we developed a hybrid fabrication strategy combining Two-photon direct laser writing (TPDLW). The pillar molds were designed using computer-aided design (CAD; SolidWorks, Dassault Systèmes). Before printing the mold, silicon substrates (University Wafer, Boston, MA) were cut to 25 mm × 25 mm. The silicon substrates were cleaned with ethanol and an air gun, followed by plasma treatment (PDC-32G, Harrick Plasma) with 10.5 W RF power for 1 minute. The substrate was then coated with 3(trimethoxysilyl)propyl acrylate (TMPA) with vapor deposition for 12 hours. The pillar molds were printed using TPDLW (Photonic Professional GT, IP-dip photoresist, Nanoscribe) with a 25X (NA = 0.8) immersion objective. The pillar master mold contained inverse cavities for a four-pillar array (1 mm x 0.6 mm) situated within a 1.12 × 1.5 mm microwell. Each pillar features a 150 µm diameter cylindrical base, a 200 µm diameter spherical cap, and a total height of 450 µm (Fig. 1B). The excess resin was removed with a propylene glycol monomethyl ether acetate (PGMEA) wash for 3-6 hours. The molds were then washed in a NOVEC 7100 (3M) bath and cured with exposure to a UV light source (5W, 389 nm, Prizmatix) for 20-50 seonds. Molds were plasma treated and subsequently fluorinated using trichloro(1H,1H,2H,2H-perfluorooctyl) silane (TPFOS, Sigma).

##### Replica Molding Pipeline

To scale fabrication of polydimethylsiloxane (PDMS, 1:10 Sylgard 184, Dow) stamps for plate casting, we first fabricated multiple PDMS replicas of the TPDLW 3D printed master mold. These replicas were then bonded to a glass slide (75 x 50 mm) with oxygen plasma in a 4x5 array configuration, treated with oxygen plasma again and then vapor coated with TPFOS for 3 hours. These slide molds were then used to cast inverse PDMS molds. These inverse PDMS molds were then vacuum cast in epoxy resin (SmoothCast 322, Smooth-On). The epoxy mold was used to fabricate multiple PDMS copies of the inverse PDMS molds. These inverse PDMS molds were also plasma treated and coated with TPFOS for 3 hours.

The 4x5 arrays of inverse stamps were then filled with drops of PDMS and glass slide (75 x 50 mm) was placed on top of the mold and clamped with binder clips. This assembly was baked at 60°C for 3 hours. The PDMS stamps cast with this process were then used downstream to assemble the PDMS-SLA 3D printed holder assemblies.

**SLA 3D Printing**

The mold holders were designed using computer-aided design (CAD; SolidWorks, Dassault Systèmes), and 3D printed with stereolithography (SLA) (Form 3+, Grey resin, Formlabs) with a 25 µm step size. The printed parts were then washed in 90% isopropyl alcohol (IPA) for 10 minutes in a Form Wash (Formlabs) and air dried. The parts were then cured in UV light at 60° C for 2 hours and heat cured overnight.

**Mold Assembly**

Multiple copies of the master mold were then fabricated in polydimethylsiloxane (PDMS, 1:10 Sylgard 184, Dow) to be used as our working PDMS stamps. To adapt these stamps for high-throughput formats, individual PDMS stamps were secured into the custom SLA 3D-printed holders using a twist-lock cap mechanism. Assembling these discrete units onto SLA 3D-printed racks yielded an 8×12 mold array with standard 9 mm spacing. This assembly was then treated with oxygen plasma  coated with TPFOS for 6 hours for further downstream use.[16,17] Because this architecture is highly modular, individual stamps can be replaced and silanized again without sacrificing the entire array, enabling the platform to be reused for more than 30 cycles with minimal maintenance.

**Plate fabrication**

To mitigate edge effects driven by uneven evaporation, we utilized 96-well plates featuring a PBS-fillable perimeter moat (Nunc Edge 2.0 96-well plate, Thermo Scientific, Cat No. 267578).[18] Within each well, PDMS devices were generated by vacuum-casting green PDMS onto the assembled 8×12 mold array, inserting the molds into the plate, and curing at 60 °C for at least 6 hours.[16,17] Once de-molded, the plate was sonicated with 70% ethanol in each well to clean the fabricated devices, air dried and stored for later use.

**Calculation and simulation of pillar effective spring constant**

Finite element modeling of the devices is implemented using COMSOL Multiphysics (Solid Mechanics Module, version 6.3). Meshes were generated using tetrahedral elements and the "extra fine" setting to determine the element size. Young's modulus of 1:10 PDMS was assumed to be 1.7 MPa based on the curing temperature of 60ºC.[19] To simulate contractile forces exerted by the tissue, we applied a 5 µN force on the surface of the spherical caps. After observing the displacement of the pillar caps ($\Delta x$), we applied Hooke's law $k \approx F/\Delta x$ to estimate the spring constant of the pillars, which was about 2 µN/µm. Our analytical estimation based on cylinders also yielded a similar result (2.07 µN/µm). (Supplementary Fig. 3)
