## Appendix-S1: WoundCompute for "A high-throughput, 3D microtissue platform for multiparametric analysis of tissue remodeling"

### APPENDIX S1 - WOUNDCOMPUTE SOFTWARE

A PREPRINT

**Quan Nguyen**
Department of Mechanical Engineering, Boston University
Boston, Massachusetts


**Anish Vasan**
Department of Biomedical Engineering and the Biological Design Center, Boston University
Boston, Massachusetts

**Emily Davis**
Department of Biomedical Engineering and the Biological Design Center, Boston University
Boston, Massachusetts

**M Çağatay Karakan**
Department of Biomedical Engineering and the Biological Design Center, Boston University
Boston, Massachusetts

**Elena Westphal**
Department of Biomedical Engineering and the Biological Design Center, Boston University
Boston, Massachusetts

**Varun Shah**
Department of Biomedical Engineering and the Biological Design Center, Boston University
Boston, Massachusetts

**Wilson Wong**
Department of Biomedical Engineering and the Biological Design Center, Boston University
Boston, Massachusetts

**Jeroen Eyckmans**
Department of Biomedical Engineering and the Biological Design Center, Boston University
Wyss Institute for Biologically Inspired Engineering, Harvard University
Boston, Massachusetts

**Emma Lejeune**
Department of Mechanical Engineering, Boston University
Boston, Massachusetts


July 19, 2026

### 1 Introduction

The WoundCompute software is a multi-purpose computational tool designed to automate the extraction and analysis of quantitative data from microtissue wound healing experiments. In these experiments, human dermal fibroblasts from neonatal foreskin (NHDFneo) or human dermal fibroblasts from adults (NHDFad) are cultured into stromal microtissues, which are then injured via nanosecond-pulsed laser ablation [Griebel et al., 2023]. The software processes high-throughput image stacks to segment both tissue and wound boundaries, and quantifies key metrics – such as wound area and major/minor wound axis lengths – while also tracking pillar positions and computing change in pillar distance from centroid of all pillars. Additionally, it assesses tissue integrity (i.e., detachment from pillars) and determines wound closure status. To validate the WoundCompute software, we compare its performance against manual analyses, demonstrating its advantages in reproducibility for wound segmentation. Then, we validate the pillar tracking algorithm by applying synthetic displacements to the microtissue experimental system, tracking the pillars with our algorithm, and confirming the accuracy of the recovered synthetic displacements. In Section 2, we detail the algorithms for segmentation, tracking, and post-processing (i.e., check tissue integrity and wound closure, compute absolute pillar displacements). Section 4 compares manual and automated analyses, highlighting the advantages of WoundCompute, while Section 5 emphasizes the modular architecture and test-driven development framework of our software. For details on the data structure used by WoundCompute and the associated experimental metadata, please refer to the "Published Microscopy Dataset" Appendix. WoundCompute thus provides a robust, standardized platform for high-throughput quantification of wound healing dynamics in engineered microtissue systems.

### 2 Methods

In this Section, we present the methods underlying our WoundCompute pipeline (Fig. 1): the automated segmentation of pillars, tissue, and wound; pillar tracking; and post-processing of the data.

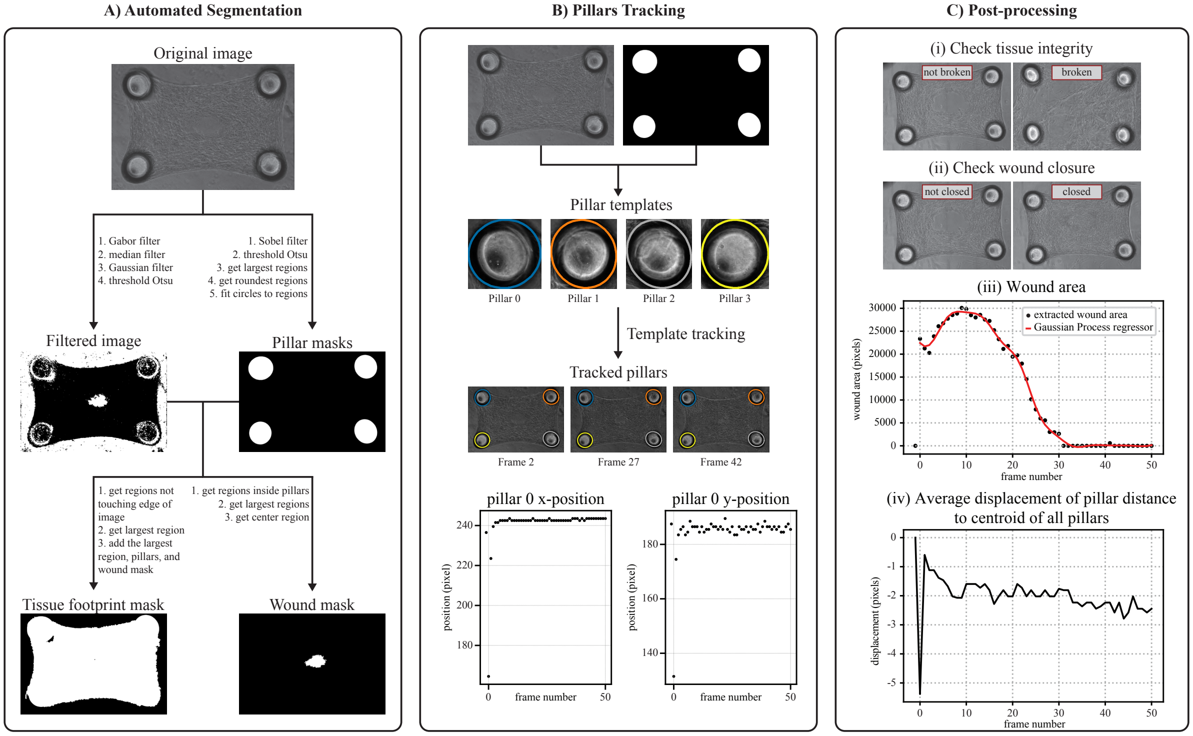


Figure 1: WoundCompute computational framework: **(A)** Automated segmentation; **(B)** Pillar tracking; and **(C)** Post-processing.

#### 2.1 Pillar segmentation

As illustrated in Fig. 2, pillars appear as dark elliptical regions against the brighter phase contrast background. Therefore, we apply an Otsu threshold [Otsu et al., 1975] to the original image to isolate the pillars from the surrounding area. The resulting binary mask is then refined through a sequence of morphological operations – including closing, inversion, erosion, and opening – proceeding in parallel tracks to produce three distinct binary images (Fig. 2A). Contours are extracted from each binary image using OpenCV [Bradski, 2000], and ellipses are fitted to all contours (Fig. 2B). From all the candidate ellipses, we remove those with an aspect ratio $< 0.50$, with an area larger than $3\%$ or smaller than $0.5\%$ of the image area. By obtaining candidate ellipses from multiple thresholding pipelines, we ensure that the true pillar regions are detected consistently, while reducing detections for spurious regions.

To match candidate ellipses with individual pillars (Fig. 2C), we cluster their centroids using K-means [Arthur and Vassilvitskii, 2007], setting the number of clusters equal to the expected number of pillars (i.e., 4 expected pillars for our experiments). This process results in 4 clusters of candidate ellipses, 1 for each pillar. Within each ellipse cluster, if the candidate ellipse centroids are spatially dispersed, we perform a second round of clustering in which the optimal number of sub-clusters is determined by maximizing the Silhouette score [Rousseeuw, 1987]. The sub-cluster containing the highest number of candidate ellipses is then selected, since true pillar regions will be represented by a greater number of candidate ellipses than spurious ones. In the case of a tie in cluster size, we prefer the sub-cluster with the highest spatial compactness (i.e., the sub-cluster with the smallest sum of distances of all points in the sub-cluster to the centroid). The final ellipse for each pillar is selected as the smallest-area ellipse within the selected sub-cluster, as the tightest fit best approximates the inner boundary of the pillar. We then expand this ellipse by a fixed buffer along both axes to cover the dark pillars. The final output are binary masks of the pillars.

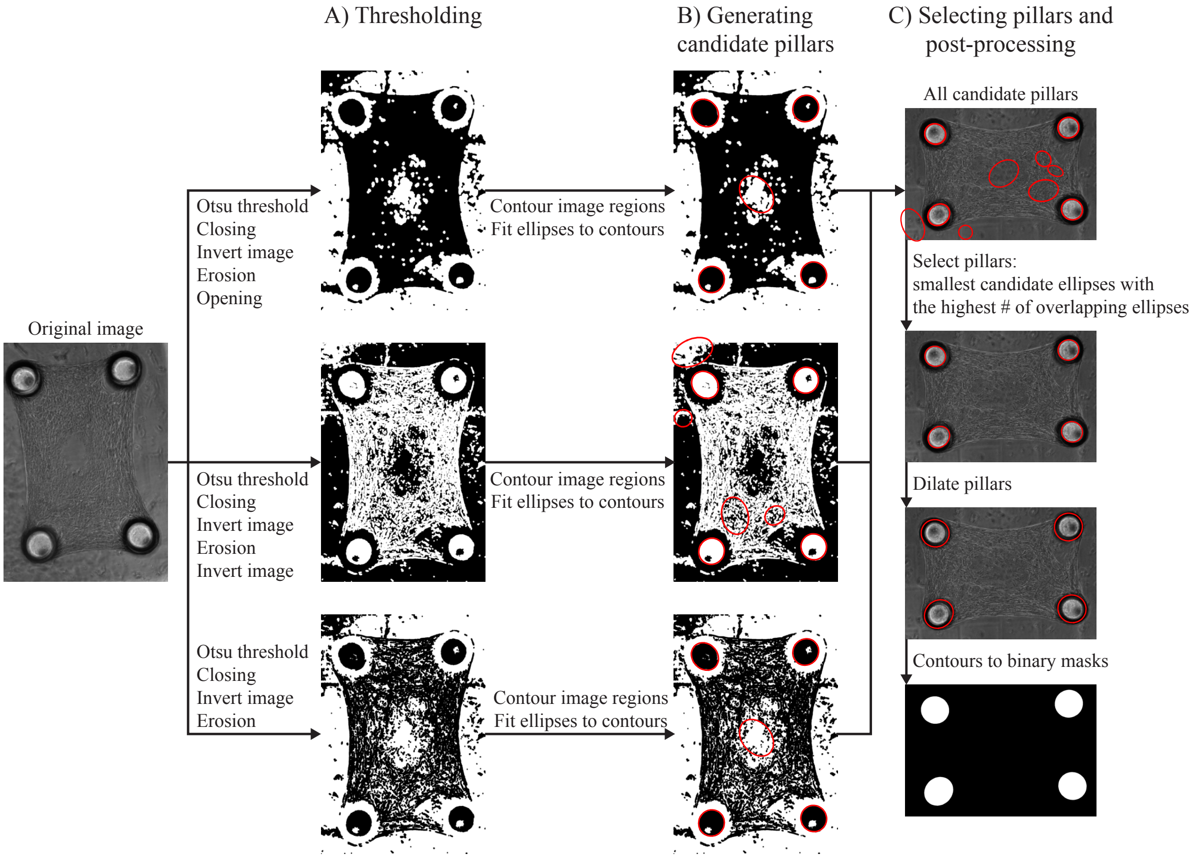


Figure 2: Pillar segmentation pipeline. **(A)** Thresholding the original images with 3 different pipelines to obtain 3 binary images. **(B)** Generating candidate pillars by obtaining the contour of all regions in an image, and fitting ellipses over the regions. **(C)** Selecting the best fit pillars and post-processing to obtain the final pillar masks.

#### 2.2 Tissue footprint segmentation

We define the tissue footprint mask as the tissue mask with the wound and pillar regions filled in (Fig. 11B). We then define tissue compaction as the change in tissue footprint area over time. This is necessary because wound area and tissue footprint are independent measures – filling in the wound and pillar regions ensures that changes in tissue footprint area reflect true compaction dynamics, independent of wound area changes. Accurate segmentation of the tissue is, however, challenging due to the low contrast between the tissue and the surrounding background, and the non-uniform texture of the tissue. When using a standard segmentation pipeline (e.g., the representative example of sequential filtering with a median filter, Gaussian filter, and Otsu thresholding), the resulting binary image contains only the pillar edges (Fig. 3A). In our exploration, we tried many other combinations of standard filters, and were not able to successfully perform segmentation. Thus, to segment the tissue and create a robust overall pipeline, we turned to the Gabor filter. The Gabor filter captures directional features of an image by convolving the image with kernels of varying orientations and frequencies (Fig. 3B) [Gabor, 1946, Granlund, 1978]. By combining multiple convolutions, the filter accentuates the tissue texture (Fig. 3C). Specifically, we choose Gabor kernels at 3 different frequencies (i.e., 0.2, 0.6, 1.0) and 17 angles – ranging from 0 rad to $\pi$ rad – for a total of 51 convolutions. The code snippet for the Gabor filter is provided in Listing 1. After combining convolutions into a single image, we apply Otsu thresholding to create a binary image of all the detected regions (Fig. 3D). The final tissue mask is obtained by selecting the largest region not intersecting the image boundary. The tissue footprint mask is then assembled by filling the wound and pillar regions into the tissue mask. Here, we include a code snippet of the Gabor filter with the help of the following Python packages – numpy [Harris et al., 2020], scipy [Virtanen et al., 2020], and scikit-image [Van der Walt et al., 2014].

```python
import numpy as np
from scipy import ndimage
from skimage.filters import gabor_kernel

def gabor_filter(
    image_array: np.ndarray,
    theta_range: int = 17,
    ff_num: int = 3,
    ff_mult: float = 0.4
    ff_base: float = 0.2
) -> np.ndarray:

    """
    Convolves a series of Gabor kernels to an input image at multiple frequencies
    and orientations. Returns the sum of the images from all Gabor convolutions.

    Parameters:
    -----------
    image_array : np.ndarray
        Input grayscale image as a 2D numpy array (height x width)
    theta_range : int, optional (default=17)
        Number of orientation angles between 0 rad and pi rad
    ff_num : int, optional (default=3)
        Number of frequency scales to apply
    ff_mult : float, optional (default=0.4)
        Frequency multiplier to increment per ff_num
    ff_base : float, optional (default=0.2)
        Base frequency for the first Gabor filter

    Returns:
    --------
    np.ndarray
        Filtered image with same dimensions as input, representing the combined
        response across all orientations and scales
    """
    # Initialize output array with same shape as input
    gabor_all = np.zeros(image_array.shape)

    # Apply filters at multiple frequencies
    for ff in range(0, ff_num):
        frequency = ff_base + ff * ff_mult  # Altering the frequency

        # Apply filters at multiple orientations
        for tt in range(0, theta_range):
            theta = tt * np.pi / (theta_range - 1) # Altering the orientation

            # Generate Gabor kernel and convolve with image
            g_kernel = gabor_kernel(frequency=frequency, theta=theta)
            filt_real = ndimage.convolve(
```

```
                image_array,
                np.real(g_kernel), # Use only real component of the Gabor kernel
                mode='reflect',  # Handle borders by mirroring
                cval=0           # Default value for points outside
            )

            # Accumulate responses across all frequencies/orientations
            gabor_all += filt_real

    return gabor_all
```

Listing 1: Gabor filter code.

##### 2.3 Wound segmentation

To segment the wound, we use the pillar masks and the filtered image (Fig. 1A), the latter generated by our Gabor filter pipeline. The wound is identified as the largest empty region fully enclosed by the pillar masks. This approach excludes smaller gaps and the empty background, ensuring the mask captures only the primary wound area. The result is a clean segmentation of the wound within the tissue boundaries (Fig. 1A).

##### 2.4 Tissue integrity check

Tissue detachment from pillars is one key mode of failure during these experiments. Since some tissues detached from the pillars during experiments, we use four conditions to check the integrity of the tissue (Fig. 4). The conditions are as follows:

1. Symmetry check (Fig. 4A) - The tissue is divided into 4 quarters using the pillar masks. For symmetry, the minimum area of the tissue in a given quarter should not exceed 0.25 of the maximum area of the tissue in a quarter. And, the ratio of the minimum area of a quarter to the mean area of all quarters should not exceed 0.60.
2. Tissue mask existence (Fig. 4B) - WoundCompute checks for the existence of a tissue mask.
3. Tissue on a single pillar check (Fig. 4C) - If the tissue area is less than $10\%$ of the total image area, the tissue might be lying on a single pillar, and is considered broken.
4. Tissue on two pillars check (Fig. 4D) - If the center of the tissue is offset from the center of the image, the tissue might be lying on two pillars, and is considered broken.

For our current experimental system, the condition to check symmetry (i.e. condition 1) captures all the broken tissues. The other conditions exist to check more complex experiments (e.g., tissues experiencing drugs, tissues located on different pillar configurations) in the future.

##### 2.5 Wound closure check

To automatically assess wound closure status, we leverage the segmented tissue mask, wound mask, and a set of conditions applied to each frame (Fig. 5). Specifically, we first rotate the tissue mask so that the longest axis of the tissue is aligned with the x-axis of the image, then we apply the same rotation matrix to the wound mask. From the rotated tissue mask, we compute the bounding box, then we scale the bounding box down by 0.25 and 0.5 – referred to as inner box and outer box respectively. We then use three conditions to check for wound closure:

1. Wound mask existence - Confirms the presence of a wound mask.
2. Wound boundary check - Ensures the entire wound lies within the outer box boundary.
3. Wound center check - Verifies that the center of the wound is inside the inner box.

If any of the above conditions fails, the wound is classified as closed.

##### 2.6 Pillar tracking

To track pillar displacements over time, we employ template matching [Brunelli, 2009] using pillar templates extracted from a reference frame (Fig. 1B). For each of the 4 pillars, a template is obtained by extracting the region defined by

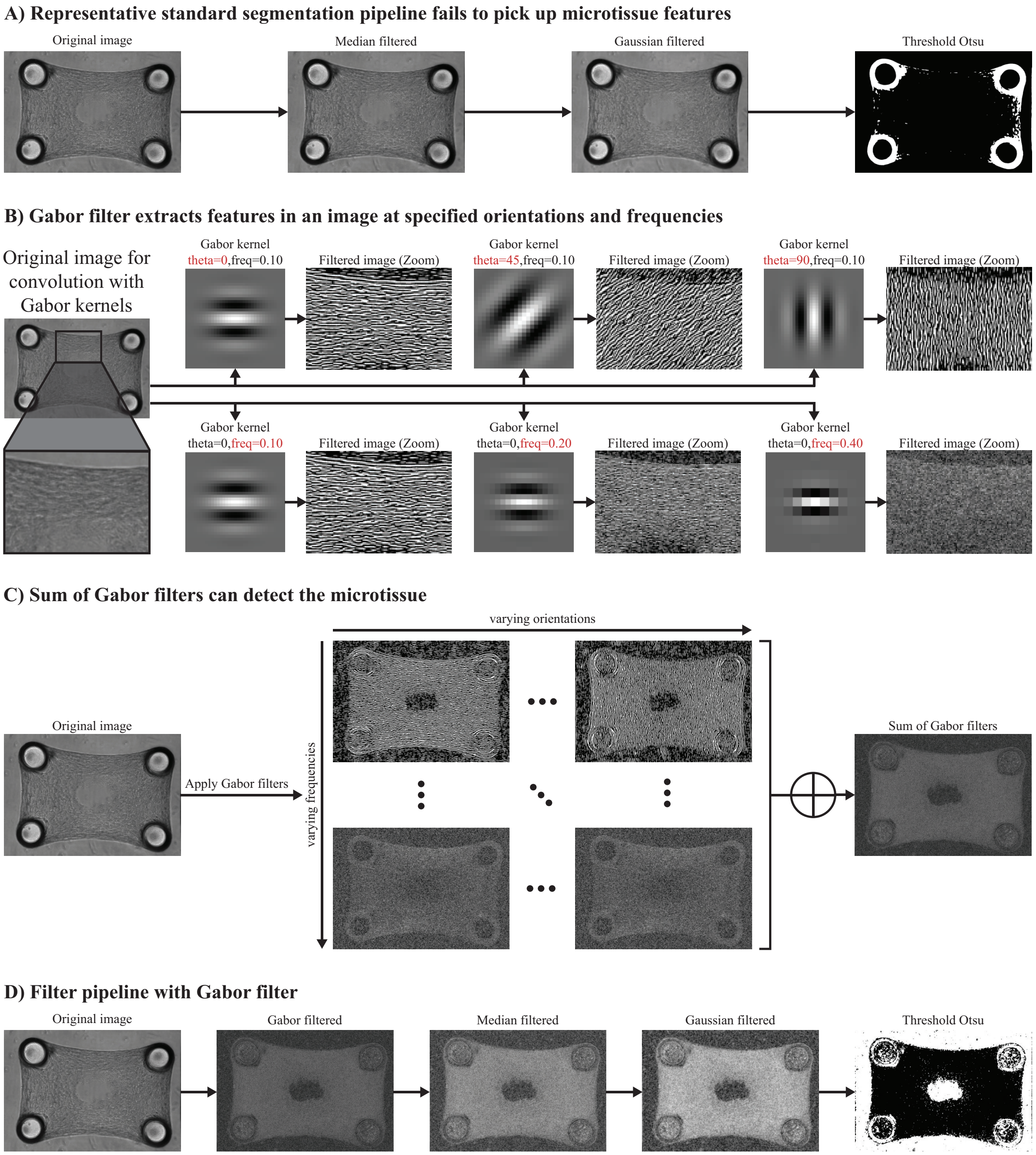


Figure 3: Functionalities and efficacy of the Gabor filter.

the pillar mask from the reference image. Tracking is then performed by sliding each template over subsequent frames and computing the normalized cross-correlation score:

$$R(x,y) = \frac{\sum_{x',y'} T(x',y') \cdot I(x+x',y+y')}{\sqrt{\sum_{x',y'} T(x',y')^2 \cdot \sum_{x',y'} I(x+x',y+y')^2}}, \quad (1)$$

where $T$ is the template, $I$ is the image, $x, y$ are the coordinates of the top-left pixel in the current window of image $I$, and $x', y'$ are the coordinates of the current pixel in template $T$. The pillar location in each frame is taken as the window with the highest score $R$, computed using the `matchTemplate` function in OpenCV [Bradski, 2000].

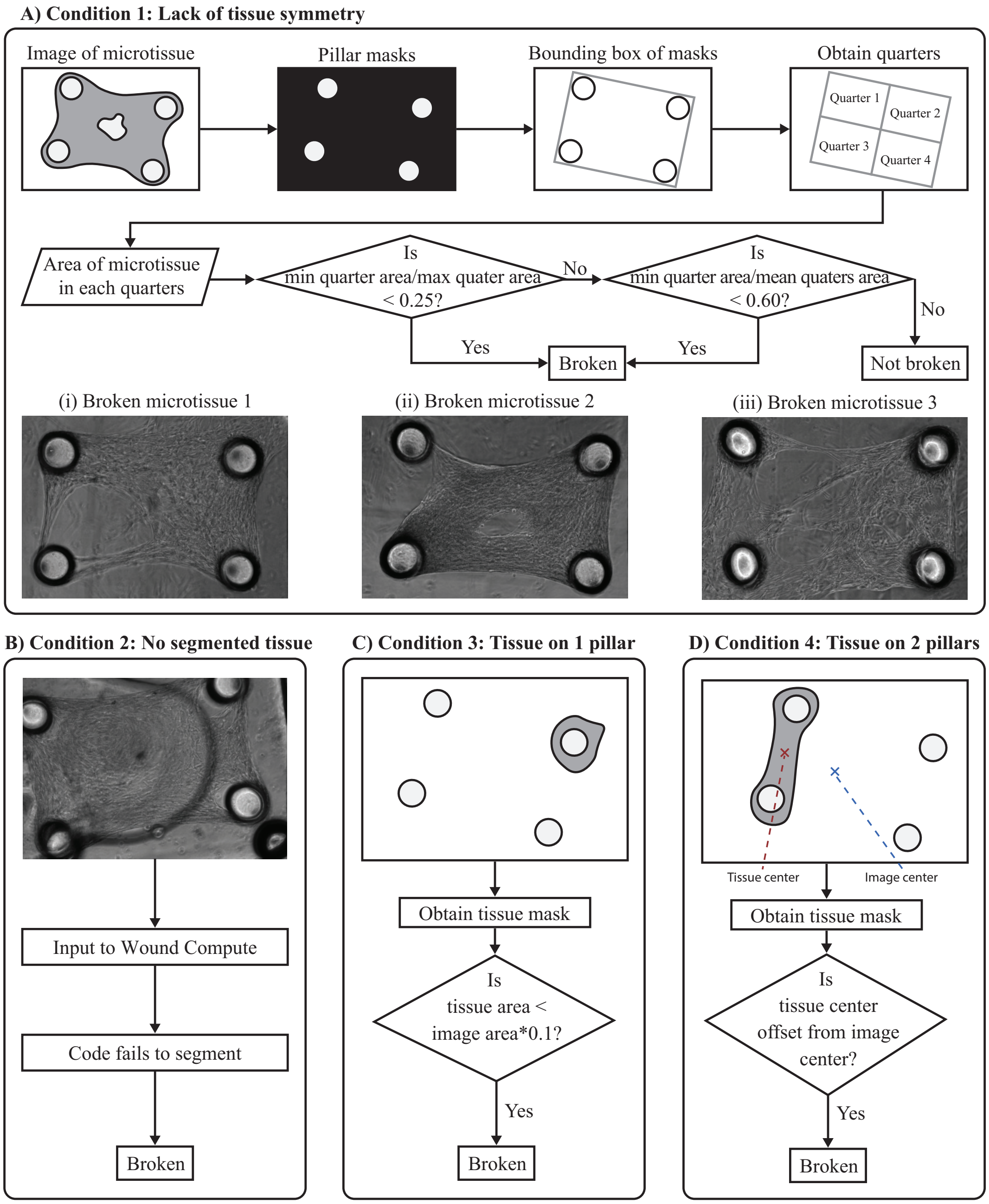


Figure 4: Schematic illustration of the conditions used to check for broken tissues.

Prior to tracking, we normalize the histogram of each frame to match that of the reference frame using histogram matching [Van der Walt et al., 2014], ensuring that correlation scores are comparable across frames. The peak correlation score of the reference frame then serves as a baseline for detecting tracking failures: if the relative absolute difference between the peak score of a given frame and the baseline exceeds 0.35, the pillar is likely moving out of the field of view. In this case, we truncate the pillar mask and template so that our algorithm only tracks the portion inside the field of view. Specifically, we identify the edge of the image closest to the pillar, cut the half of the mask (and template) closest to that edge, and retain the half of the mask closer to the center of the image. Tracking is then repeated with the truncated template to recover the pillar location.

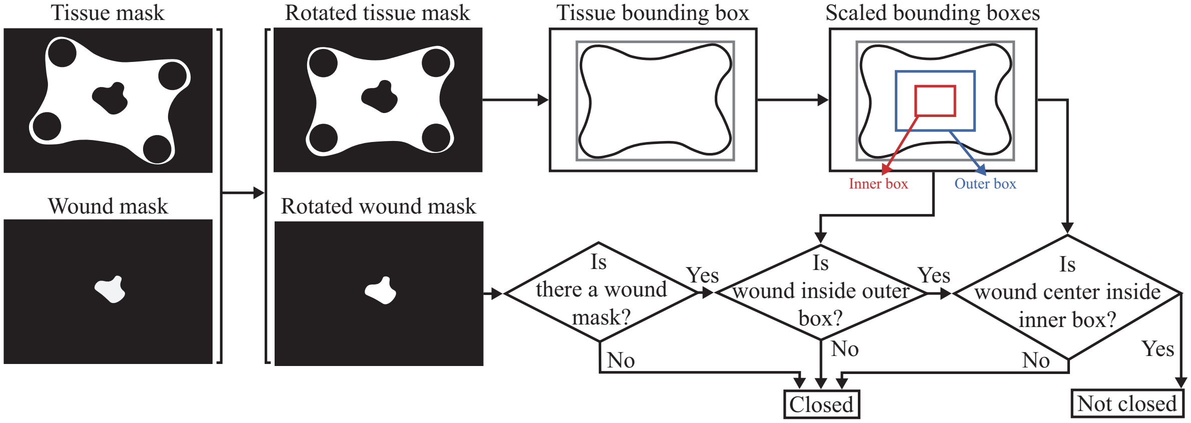


Figure 5: Schematic illustration of the pipeline to check wound closure.

##### 2.7 Change in pillar distance from centroid of all pillars

To obtain the change in pillar force while accounting for global drift, we first compute the change in distance of each pillar to the centroid of all pillars over time (Fig. 6). Since global drift shifts all pillars equally, it does not affect the distance of each pillar to their shared centroid, making this metric robust to drift. At each frame, the centroid of all pillars is computed from the current pillar positions as:

$$\bar{x}(t) = \frac{1}{N}\sum_{i=1}^{N} x_i(t), \quad \bar{y}(t) = \frac{1}{N}\sum_{i=1}^{N} y_i(t), \tag{2}$$

where $x_i(t)$ and $y_i(t)$ are the position of pillar $i$ at frame $t$ and $N$ is the number of pillars. The distance of each pillar center to this centroid is then calculated as:

$$d_i(t) = \sqrt{(x_i(t) - \bar{x}(t))^2 + (y_i(t) - \bar{y}(t))^2}. \tag{3}$$

The change in distance of each pillar from the centroid (Fig. 6B) is computed relative to the reference frame:

$$\Delta d_i(t) = d_i(t) - d_i(0), \tag{4}$$

where $d_i(0)$ is the distance of pillar $i$ to the centroid at time 0, immediately after injury. The average change in distance across all pillars at each frame (Fig. 6C) is also reported:

$$\overline{\Delta d}(t) = \frac{1}{N}\sum_{i=1}^{N} \Delta d_i(t). \tag{5}$$

##### 2.8 Smooth noise in wound and tissue footprint area over time with Gaussian Process Regression

During the area computation for the wound and tissue footprint masks (Fig. 10A, Fig. 11A), we observe small fluctuations in the area over time graphs. Since the fluctuations occur due to noises in our experiments, we mitigate these fluctuations by applying Gaussian Process Regression (GPR), a non-parametric Bayesian approach that models the data (i.e., wound and tissue footprint area) as a distribution over functions [Williams and Rasmussen, 2006]. GPR assumes that the relative distance measurements $y(t)$ at time points $t$ are sampled from a multivariate Gaussian distribution:

$$y(t) \sim \mathcal{GP}(m(t), k(t, t')), \tag{6}$$

where $m(t)$ is the mean function, and $k(t, t')$ the covariance (kernel) function that defines the correlation between time points $t$ and $t'$. In our case, we use the radial basis function kernel [Williams and Rasmussen, 2006], which allows us to smoothly interpolate the wound and tissue footprint area through the noises:

$$k(t, t') = e^{-\gamma||t-t'||^2}, \tag{7}$$

where $\gamma = \frac{1}{2\sigma^2}$ is the signal variance.

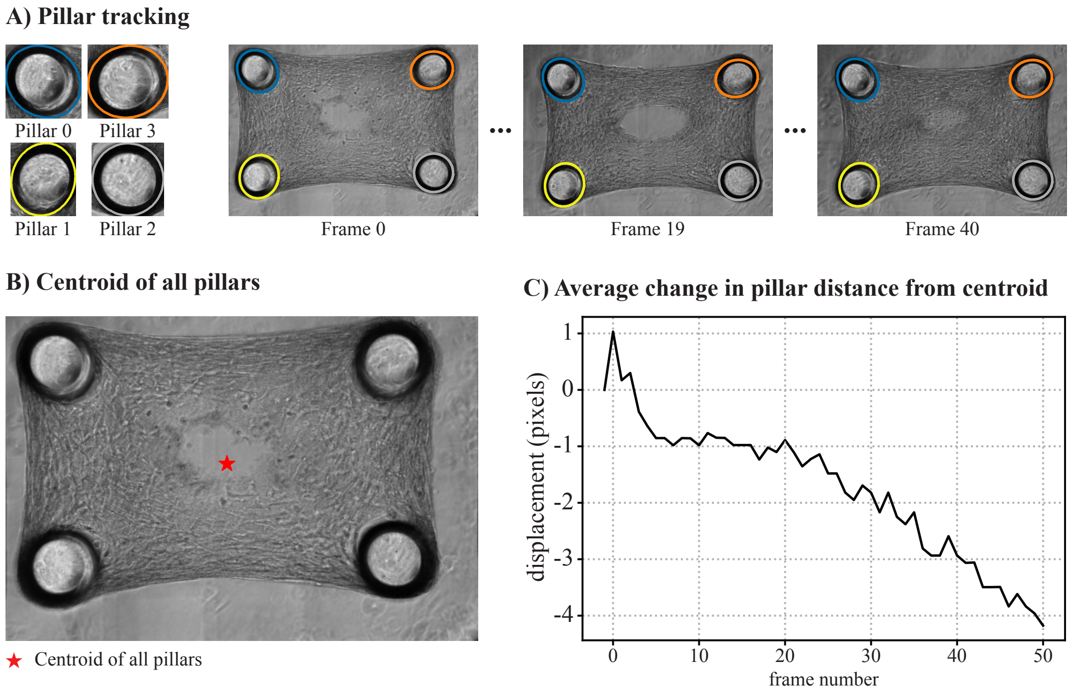


Figure 6: Pillar tracking post-processing: **(A)** template tracking of pillars; **(B)** centroid of all 4 pillars; **(C)** average change in pillar distance from centroid.

##### 2.8.1 Processing low quality frames

Certain frames within an experiment may suffer from isolated quality issues, such as being out-of-focus or exhibiting sudden illumination changes. These frames are identified manually and their indices recorded in the `.yaml` file for each experiment – for example, if frames 0 and 1 are out-of-focus, the user sets `low_quality_file_inds = [0, 1]`. Since imaging conditions in a 96-well experiment affect all wells simultaneously, the WoundCompute GUI also allows low quality frame indices to be specified at the experiment level. These recorded indices ensure that no low quality frame is selected as the reference frame for tracking; instead, the reference frame is automatically set to the closest subsequent frame not marked as low quality. Low quality frames are otherwise retained and processed normally by WoundCompute, as the quality issues do not always affect all aspects of the analysis. However, in the final analysis, frames for which the wound margin cannot be reliably delineated are manually excluded from conclusions drawn from wound area measurements. The specific frames excluded are documented in the Published Microscopy Dataset appendix.

#### 3 Results - Representative Examples

This section presents representative outputs generated by WoundCompute to illustrate its analytical capabilities; a summary of each output type and its corresponding figure is provided in Table 1.

Table 1: Summary of WoundCompute representative outputs.

| **Output** | **Figure** |
|---|---|
| Wound segmentation | Fig. 7A |
| Wound closure status | Fig. 7A |
| Microtissue integrity assessment | Fig. 7B |
| Temporal evolution of wound area | Fig. 10A |
| Temporal evolution of tissue footprint area | Fig. 11A |
| Average change in pillar distance from centroid | Fig. 6C |

First, we take a look at the main segmentation and tracking results from WoundCompute. In Figure 7, we display the segmentation results overlaid on phase contrast images, along with annotations for wound closure and tissue integrity.

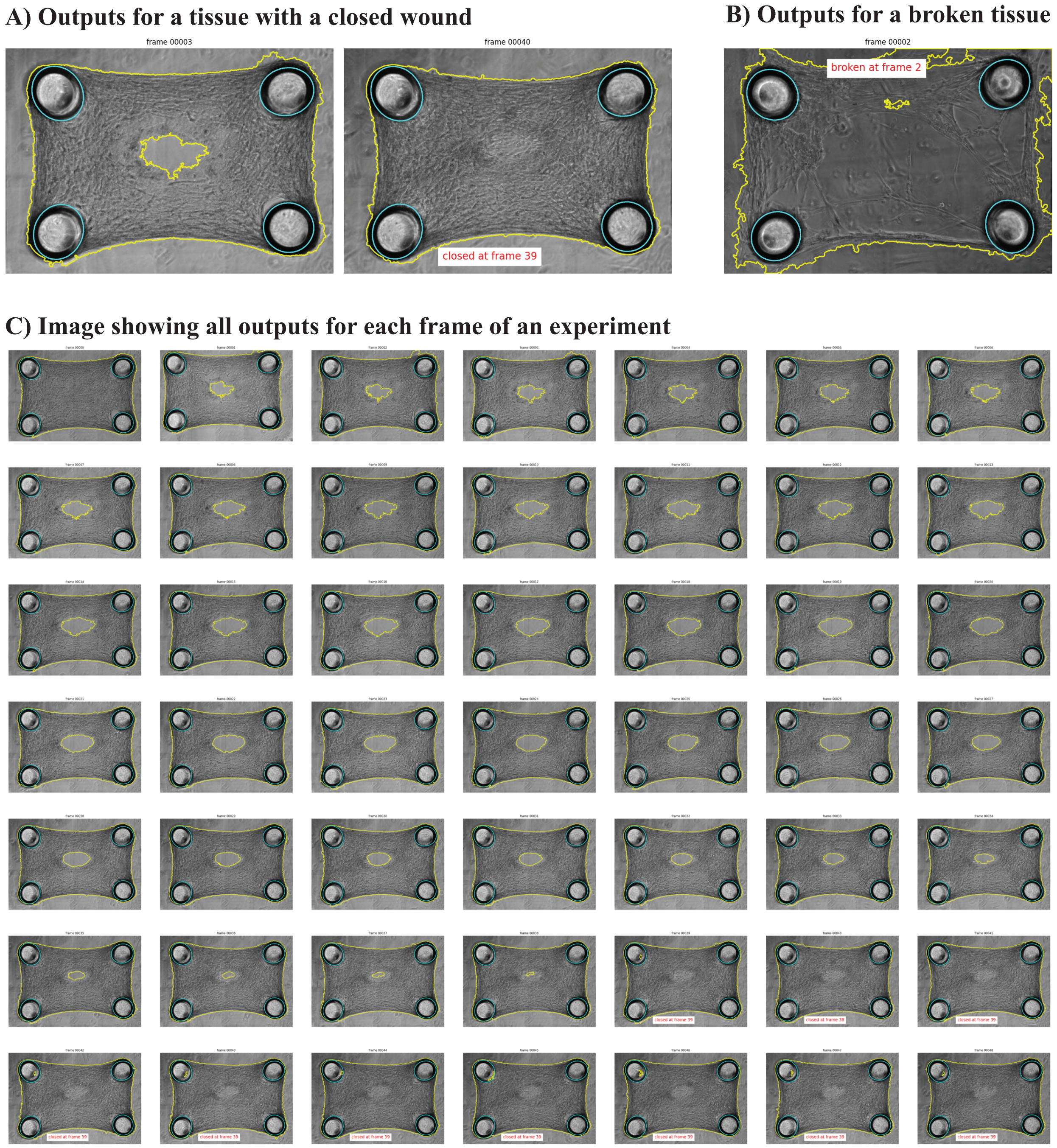


Figure 7: Segmentation results from WoundCompute: **(A)** Segmented wound, tissue, and closure status for an intact microtissue - Viability 1 well B04; **(B)** Representative broken status for a sample - Viability 1 well H04; **(C)** Segmentation results for all frames in an experiment (Viability 1 well B04), available as both .png and .gif.

For each frame, the wound contour, the tissue contour, and the pillar contours are superimposed on the raw image (Fig. 7A). If the wound is considered closed, a text box appears indicating the corresponding frame number (e.g., 'closed in frame [frame number]'). Similarly, if WoundCompute detects a structural break in the tissue, a warning message (e.g., broken at frame [frame number]) appears at the top of the image (Fig. 7B). Given the high-throughput nature of our platform, we provide quick per-sample overviews displaying all segmented frames in a single image (segment_ph1/visualizations/ph1_contour_all_[sample name].png), and animated .gif files of all segmented frames, located in the same directory.

From the segmented wound masks, the wound area is calculated as the sum of the pixels in the mask for each time point and plotted as a function of time (Fig. 10A). While the results are generally smooth due to the robustness of our algorithm, minor fluctuations may still arise. These are often attributed to background artifacts, such as floating or blurred cells, which may be occasionally misclassified as wound boundaries. Additionally, because images are captured every 30 minutes, wound area changes can appear more abrupt than expected. To address this, we apply GPR to smooth the wound area graph. The wound segmentation results for the initial time (frame 0), time at peak wound size (frame 19), time at half wound closure (frame 30), and time at wound closure (frame 40) are shown in Fig. 10B. The corresponding wound area plots are saved in the folder [sample name]/segment_ph1/.

The tissue footprint area is computed as the area covers by the tissue, wound and pillars (Fig. 11A). Here, we observe some small fluctuations between consecutive frames. The source of these fluctuations could come from cells floating near the edge of the tissue, and the less clear margin between the tissue and the background in some cases. We again smooth the tissue footprint area using GPR to clear up some noises.

In Fig. 6, we show the pillar tracking and post-processing results. The pillar labels (e.g., p0, p1, p2, p3) and the corresponding pillar templates are visualized in Fig. 6A. The tracked position of each pillar is displayed in Fig. 6B. Although the pillars displace minimally throughout the experiments, we still observe a trend in the pillar positions. To analyze only the pillar deflections and exclude any rigid motion caused by the background, we follow the procedure in Section 2.7, and plot the average change in pillar distance from centroid in Fig. 6C. We observe that the absolute pillar displacements increase linearly over time. All pillar-related outputs, including label visualizations and absolute pillar displacements plots, are located in [sample name]/track_pillars_ph1/.

##### 3.1 Run time

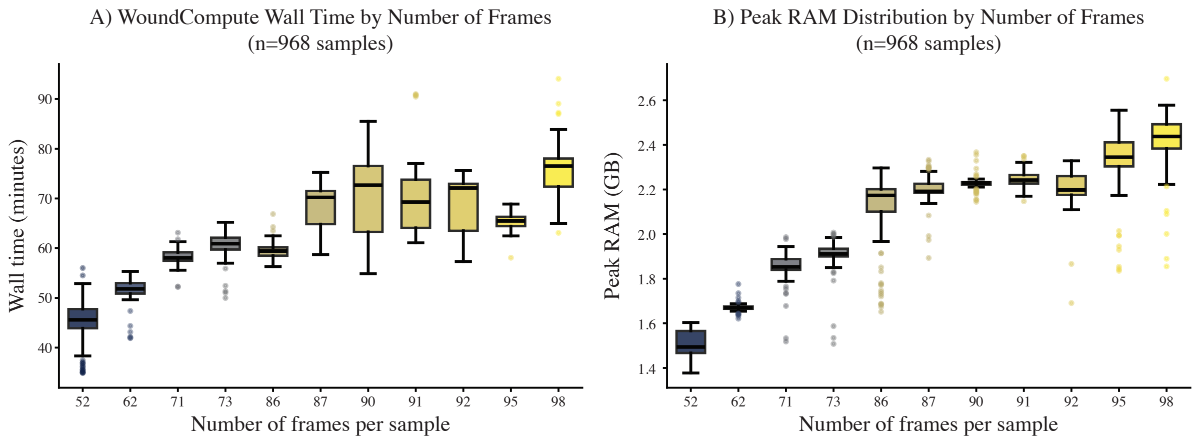


Figure 8: Computational performance of WoundCompute across 968 successfully processed samples, colored by number of frames per sample. **(A)** Wall time and **(B)** peak RAM usage as a function of the number of frames per well. Both metrics scale approximately linearly with frame count. Benchmarks were performed on Intel Xeon Gold 6242 nodes (2.80 GHz) on the Boston University Shared Computing Cluster (SCC), with one CPU core allocated per well and an image resolution of $1608 \times 1104$ pixels.

We benchmarked WoundCompute on the Boston University Shared Computing Cluster (SCC), running all 1062 wells in parallel, where each well was processed as an independent job. All jobs were pinned to Intel Xeon Gold 6242 nodes (2.80 GHz, 32 cores, 187 GB RAM) to ensure consistent hardware across all samples. Of the 1062 wells, 968 completed successfully. The remaining 94 were excluded due to experimental image quality issues, including imaging artifacts, broken tissues, and pillars completely out of frame. All benchmarks were performed on images with a resolution of $1608 \times 1104$ pixels; run time is expected to scale with image resolution, though this was not characterized as all experiments in this dataset share the same resolution.

Figure 8 summarizes wall time and peak RAM usage as a function of the number of frames per well. Both metrics scale approximately linearly with frame count ($R^2 = 0.77$), with wall time following the relationship $t \approx 0.59f + 15.1$ minutes, where $f$ is the number of frames. Across all successful wells, the median wall time was 61.6 min (interquartile range [IQR]: $53.2 - 71.0$ min) and the median peak RAM usage was 2.17 GB (IQR: $1.68 - 2.26$ GB, max: 2.70 GB).

Each well was processed in a single-threaded manner (1 CPU slot per job), and the computational requirements are well within the capabilities of a modern laptop or desktop. Users without access to a computing cluster can process wells sequentially on a local machine; summing the wall times across all 968 successful wells gives a total of approximately 995 hours, which represents the expected sequential processing time on hardware with comparable single-core performance. For faster local processing, WoundComputeGUI includes built-in support for parallel processing across available CPU cores. To demonstrate the reasonable run time achievable on personal devices, a test of 88 wells with 56 frames each was run in parallel on a laptop equipped with an AMD Ryzen 9 5900X processor (12 cores, 3.70 GHz) and 64 GB DDR4 RAM running Windows 11. The processing was completed in $\sim$3 hours 30 minutes, with a mean CPU utilization of 95%.

### 4 Validation

Manual segmentation results from human annotators are typically used as the ground-truth for validation, or as the ground-truth for training machine learning models. This makes manual segmentation the current state-of-the-art method. However, manual segmentation can have large variations between human annotators [Mohammadzadeh and Lejeune, 2025], especially as the contrast between the background and the objects decreases. And, manual segmentation is time consuming, which increases fatigue in human annotators, and reduces the quality of the segmentation along with the total number of images they can process. Due to these limitations, we are interested in assessing the quality of the manual labels in conjunction with validating our WoundCompute software based on them. The remainder of this section is as follows: (1) we compare the manual segmentation results (tissue footprint segmentation, wound segmentation, and pillar displacements) to see the variability; (2) we compare the WoundCompute results (tissue segmentation, wound segmentation) to the manual results; (3) finally, we come up with a synthetic displacement method to validate the pillar tracking results of WoundCompute due to the demonstrated limitations of manual segmentation.

#### 4.1 Variability between manual annotators - Wound segmentation

Given a stack of experimental images, we ask 4 manual annotators to segment the wound, the tissue, and the pillars based on consistent instructions and heuristics. In Fig. 9, we observe variability between the manual annotators despite our efforts to maintain consistency. In our experiments, the initial wound has a highly irregular shape, with color deviations only slightly from the gray background. These complicated areas are especially difficult to reproduce for manual annotators (Fig. 9A(iv)).

#### 4.2 Variability between manual annotators - Tissue segmentation

For tissue segmentation, we again provide the manual annotators the same stacks of images and heuristics. Due to the higher contrast between the tissue and the background, as well as the simpler shape of the tissue, tissue segmentation results are much more consistent between manual annotators. In Fig. 9, we observe that the IOU scores between the manual annotators are all above 0.95, indicating high overlap and similarity between annotators. In these simpler cases, where the margins between the objects and the background are clearer, manual segmentation becomes a reliable means to obtain the object masks.

#### 4.3 Variability between manual annotators - Pillar tracking

To track the pillars between frames, the manual annotators are asked to mask the pillars with circles (Fig. 12A). Although the dark circular pillars are visually distinct from the light gray background and the tissue, the task of pillar tracking is still challenging due to the small pillar displacements in our experiments (i.e., displacements ranging from 0 to 15 pixels in our 1104 by 1608 images). Hence, even small inconsistencies are impactful. In Fig. 12A, the positions of the red pillar masks relative to the pillars are offset by 1–3 pixels between frame 4 and frame 5. These masking inconsistencies, occurring over multiple frames, cause the high variability between manual annotators in the computed absolute pillar displacement (Fig. 12B). Therefore, instead of using manual pillar tracking as our ground truth to

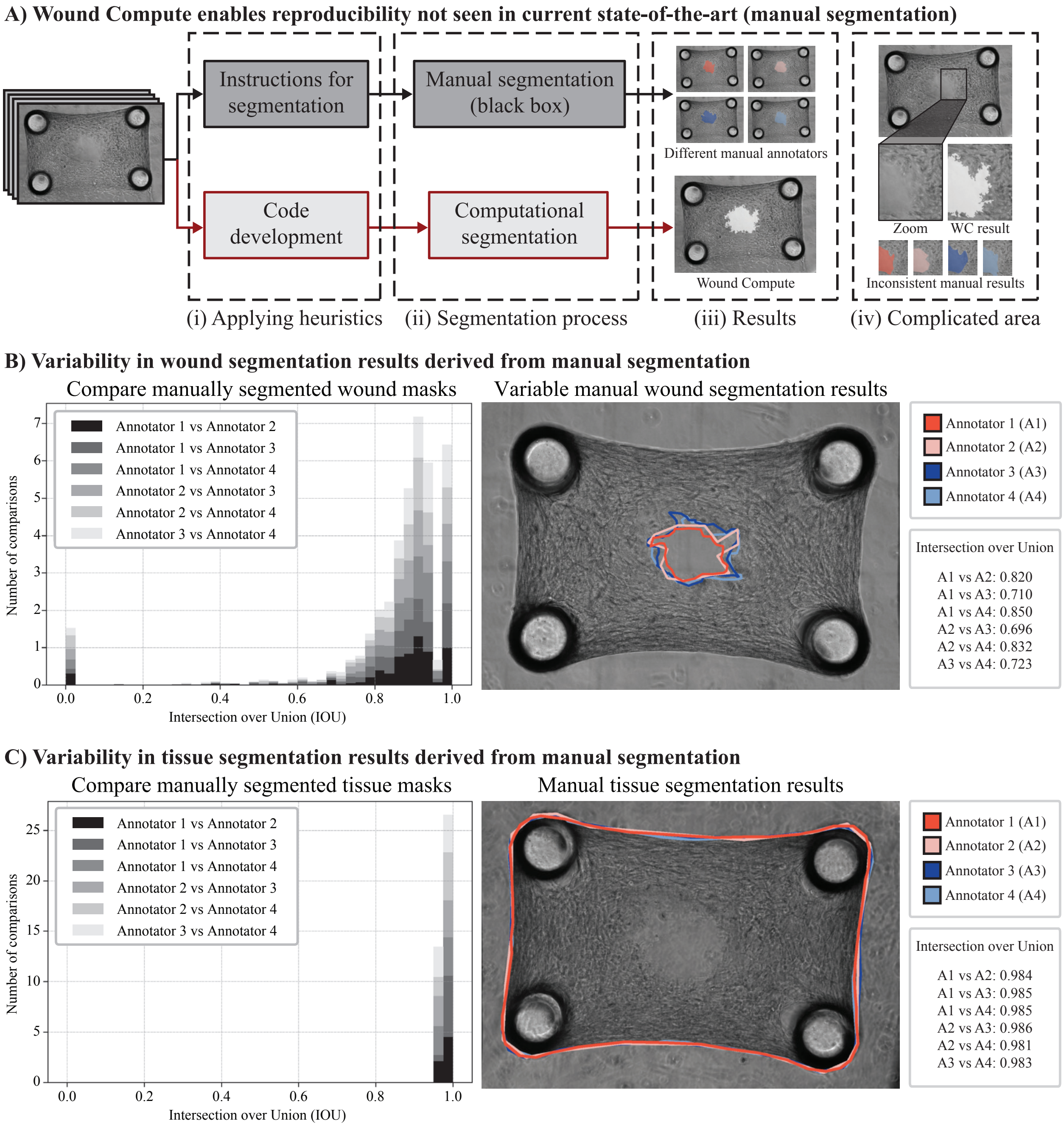


Figure 9: Reproducibility in manual wound segmentation: **(A)** WoundCompute results are reproducible; **(B)** Manual wound segmentation results vary between annotators; **(C)** Manual tissue segmentation results are consistent between annotators.

validate WoundCompute, we develop a synthetic displacement strategy to ensure the reliability of our software in Section 4.5.

##### 4.4 WoundCompute versus manual annotators

While manual segmentation is prone to reproducibility issues stemming from inter-annotator variability, subjective boundary interpretation, and annotator fatigue, automated segmentation with WoundCompute enables consistent results across a high number of images. In automated segmentation, instead of providing written or verbal instructions, we

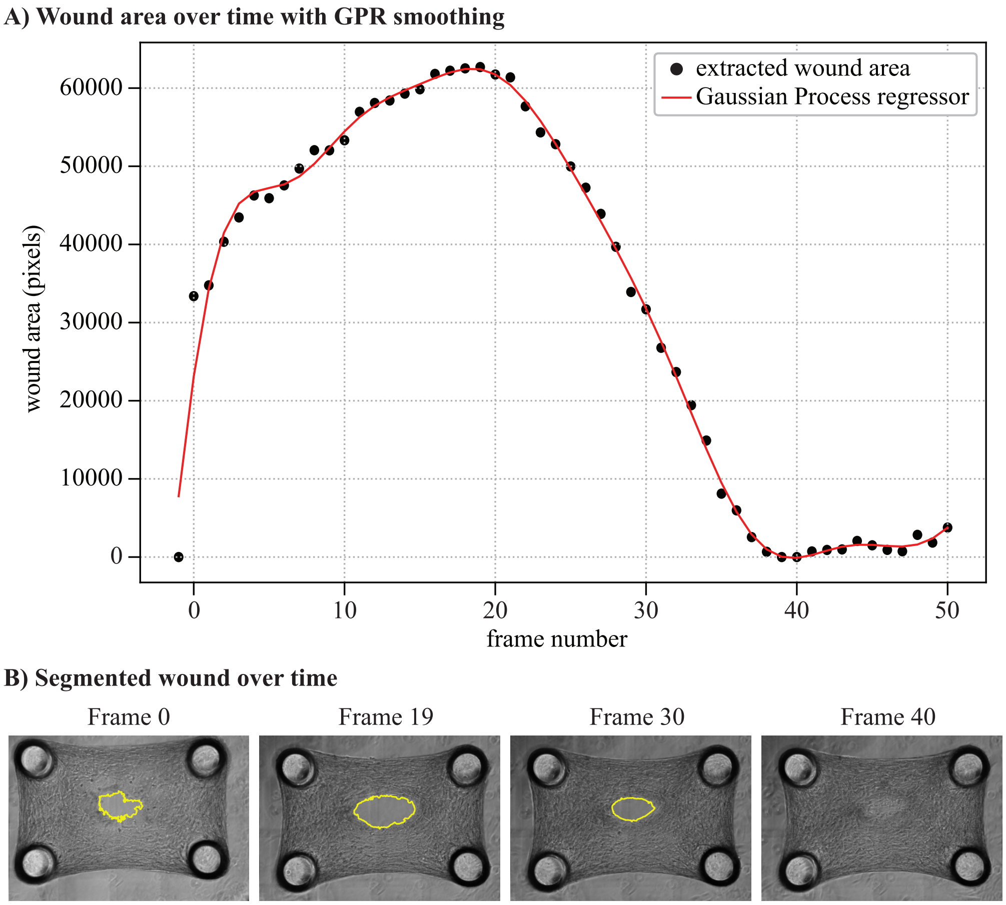


Figure 10: Results for wound segmentation over time (Viability 1 sample 22). **(A)** Wound area over time with Gaussian Process Regression smoothing. **(B)** Segmented wound over time.

apply heuristics directly to our codes. Then, we process the images via our software for analysis. Using the given heuristics, WoundCompute assesses every single pixel across all images in our high-throughput system, which provides results much more consistent than that of manual annotators. Despite the variability in the shape of the manual wound segmentation (Fig. 9B), the resulting manual wound areas are consistent. Hence, we can compare the wound areas from manual annotators to WoundCompute for validation. In Fig. 13A, we observe that WoundCompute wound areas are close to that of the manual annotators, which means that the wound masks segmented by our software are plausible. Fig. 13C shows a wound segmentation results between WoundCompute and Annotator 1 – similar masks with WoundCompute being more detailed. While most of the data points in Fig. 13A are linearly correlated, there are some data points that show a significant difference between WoundCompute and manual annotators (i.e., points with black edges showing disparity in wound closure). At these points, WoundCompute has determined that the wound is closed, while the manual annotators think that the wound is still opened. This discrepancy occurs because the annotators segment every 5 frames, so when the wound closes and reopens, the annotators might miss the previous frames closure. We note that in this dataset, when the wounds close and reopen, they always close again.

To validate the WoundCompute tissue footprint segmentation results, we compare the tissue footprint mask areas from WoundCompute to manual annotators (Fig. 13B). The tissue footprint mask (from Section 2.2) covers the area of the tissue, the wound, and the micropillars. Since the wound area changes significantly over time, the use of the tissue footprint mask removes the effects of the wound area in our comparison. Here, we find that the tissue footprint areas are generally consistent between our software and WoundCompute. For tissue footprint segmentation, the segmentation masks between WoundCompute and manual annotators are similar due to the distinct tissue boundary relative to the background (Fig. 13D).

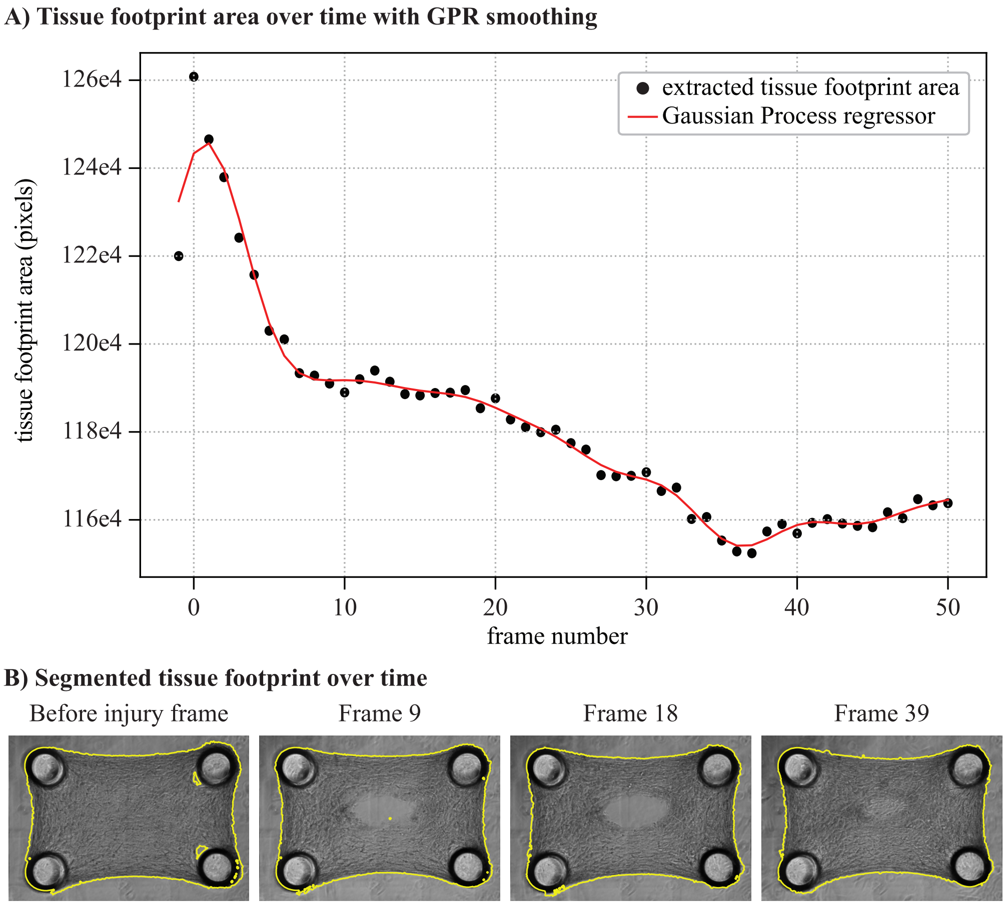


Figure 11: Results for tissue segmentation over time (Viability 1 sample 22). **(A)** Tissue area over time with Gaussian Process Regression smoothing. **(B)** Segmented tissue over time.

##### 4.5 Pillar tracking validation with synthetic displacements

Due to the inconsistency in manual pillar tracking (Section 4.3), we cannot rely on manual data to validate WoundCompute. Instead, we develop a synthetic displacement validation pipeline to ensure the reliability and reproducibility of our pillar tracking algorithm (Fig. 14). Specifically, we apply controlled synthetic displacements to the pillars, and assess the ability of our algorithm to track and recover these synthetic displacements. Because the pillars are physically connected to the tissue, any synthetic displacement applied to the pillars must also be applied to the entire system – including the tissue and wound – to preserve the relative motion observed experimentally. For each frame, we generated uniform random synthetic displacements within a predefined range, ensuring the displaced tissue remained within the field of view (Fig. 14A). The pillars were then tracked using a template matching algorithm (Section 2.6). Notably, the tracked displacements comprise both the applied synthetic displacements and the inherent experimental displacements caused by tissue motion and background movement:

$$\Delta x^{template\,tracked} = \Delta x^{synthetic} + \Delta x^{experimental}, \tag{8}$$

$$\Delta y^{template\,tracked} = \Delta y^{synthetic} + \Delta y^{experimental}, \tag{9}$$

To evaluate tracking accuracy, we compute the recovered synthetic displacements by subtracting the experimental displacements from the tracked displacements:

$$\Delta x^{synthetic} = \Delta x^{template\,tracked} - \Delta x^{experimental}, \tag{10}$$

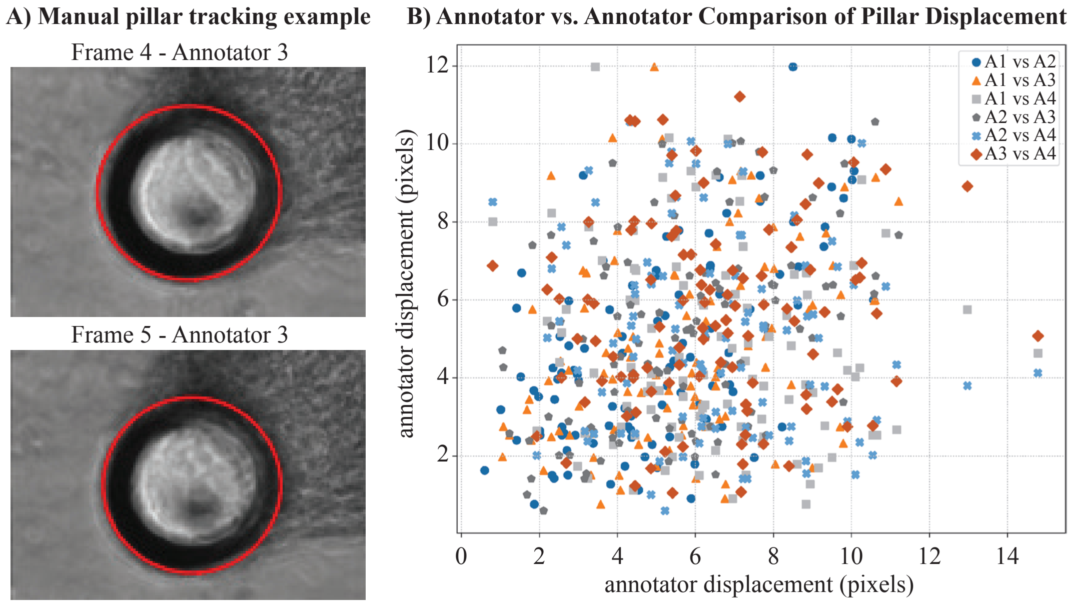


Figure 12: Pillar displacement results between manual annotators: **(A)** Manual pillar tracking example; **(B)** Annotator vs. annotator comparison of pillar displacement. We observe high variability between annotators due to the relatively small displacements (1 – 12 pixels in a 1608x1104 image) of the pillars.

$$\Delta y^{synthetic} = \Delta y^{template\,tracked} - \Delta y^{experimental}, \tag{11}$$

If the experimental displacements are correctly tracked, the recovered synthetic displacements should match the applied synthetic displacements. As shown in Fig. 14C, we observed a strong linear correlation between the recovered and applied synthetic displacements, confirming the accuracy of our pillar tracking algorithm. To further study the effect of how noise affects our tracking results, we apply Gaussian filters with varying noise level to our image stacks before tracking (Fig. 15A). In Fig. 15B, as the noise level increases, the recovered synthetic displacements gradually become less similar to the actual synthetic displacements. However, even with the added noise, the tracking results are still reasonably accurate due to the contrast between the pillars and the background.

#### 5 Future extensions of this work

WoundCompute employs a modular architecture, partitioning key workflows such as image preprocessing, segmentation, and quantitative analysis into self-contained components. This design allows for targeted updates (e.g., replacing the Gabor filter based segmentation pipeline with a machine learning approach) without systemic disruption. To ensure reliability, we adopt test-driven development (TDD), with unit tests for individual modules and integration tests for cross-component behavior. The modular approach simplifies maintenance by isolating dependencies, facilitates collaborative development by enabling parallel work on separate components, and supports scalability when extending functionality. Combined with TDD, this reduces unintended side effects during iterative refinement of WoundCompute, particularly when adapting to challenging datasets or new methodologies. Future work on WoundCompute focuses on 2 main areas: (1) expanding the software to capture more biologically relevant behaviors (e.g., full-field displacements of microtissue), and (2) enhancing WoundCompute robustness against noise and on different imaging modalities. By grounding future development in a principled software architecture, WoundCompute is well-positioned to grow alongside advances in microtissue engineering and high-throughput biological imaging.

#### 6 Data and Code Availability

WoundCompute source code is available at `https://github.com/elejeune11/woundcompute` (archived on Zenodo, `https://doi.org/10.5281/zenodo.19830254`), and a graphical interface for non-computational users is available at `https://github.com/quan4444/woundcomputeGUI`. The accompanying microscopy dataset is detailed in Appendix S2, with the full dataset publicly available through the BioImage Archive at

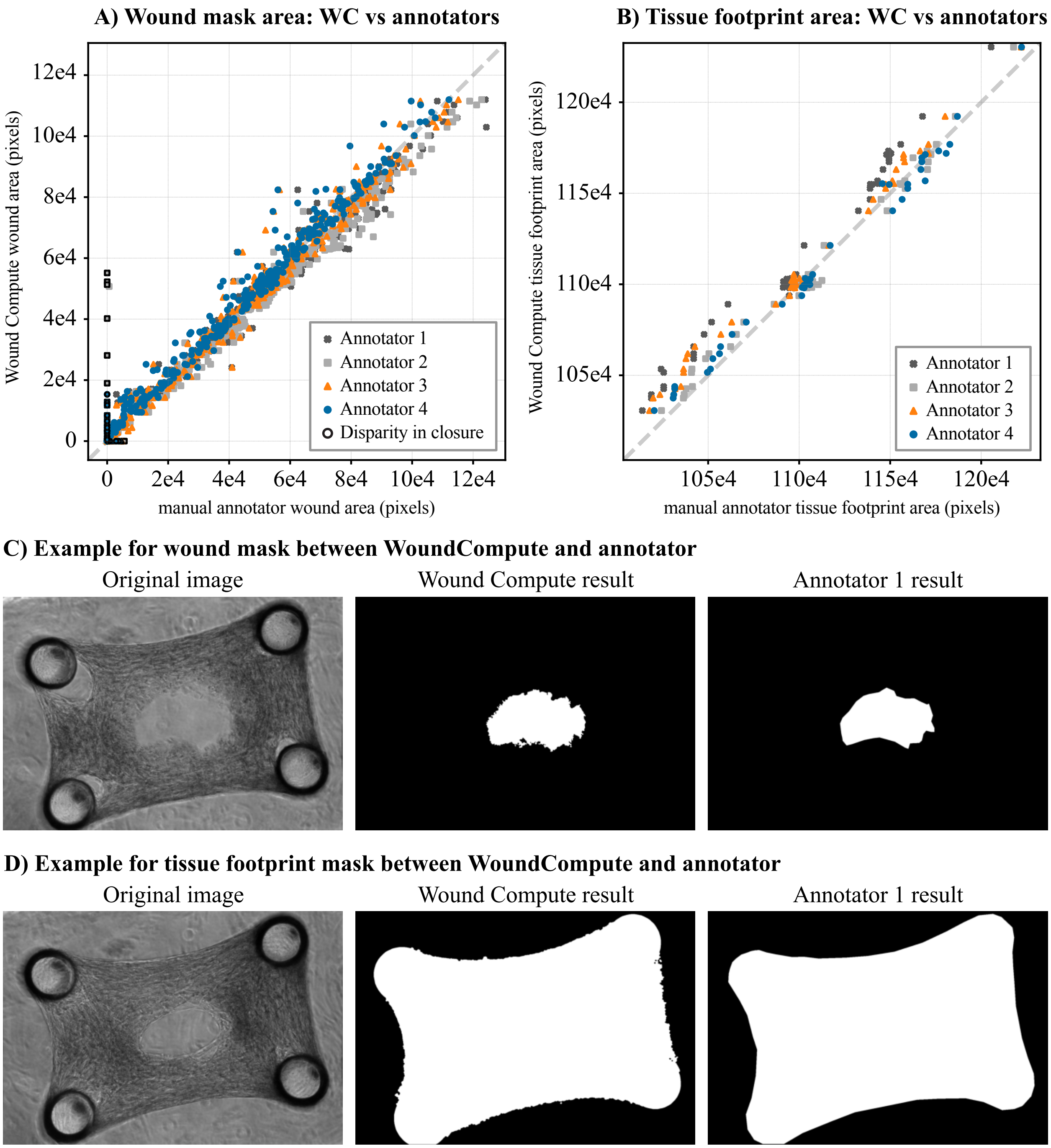


Figure 13: Comparing segmented tissue and wound area between WoundCompute and manual annotators: **(A)** Wound mask area between WoundCompute and manual annotators; **(B)** Tissue footprint mask area between WoundCompute and manual annotators; **(C)** Wound mask between WoundCompute and Annotator 1; **(D)** Tissue footprint mask between WoundCompute and annotator.

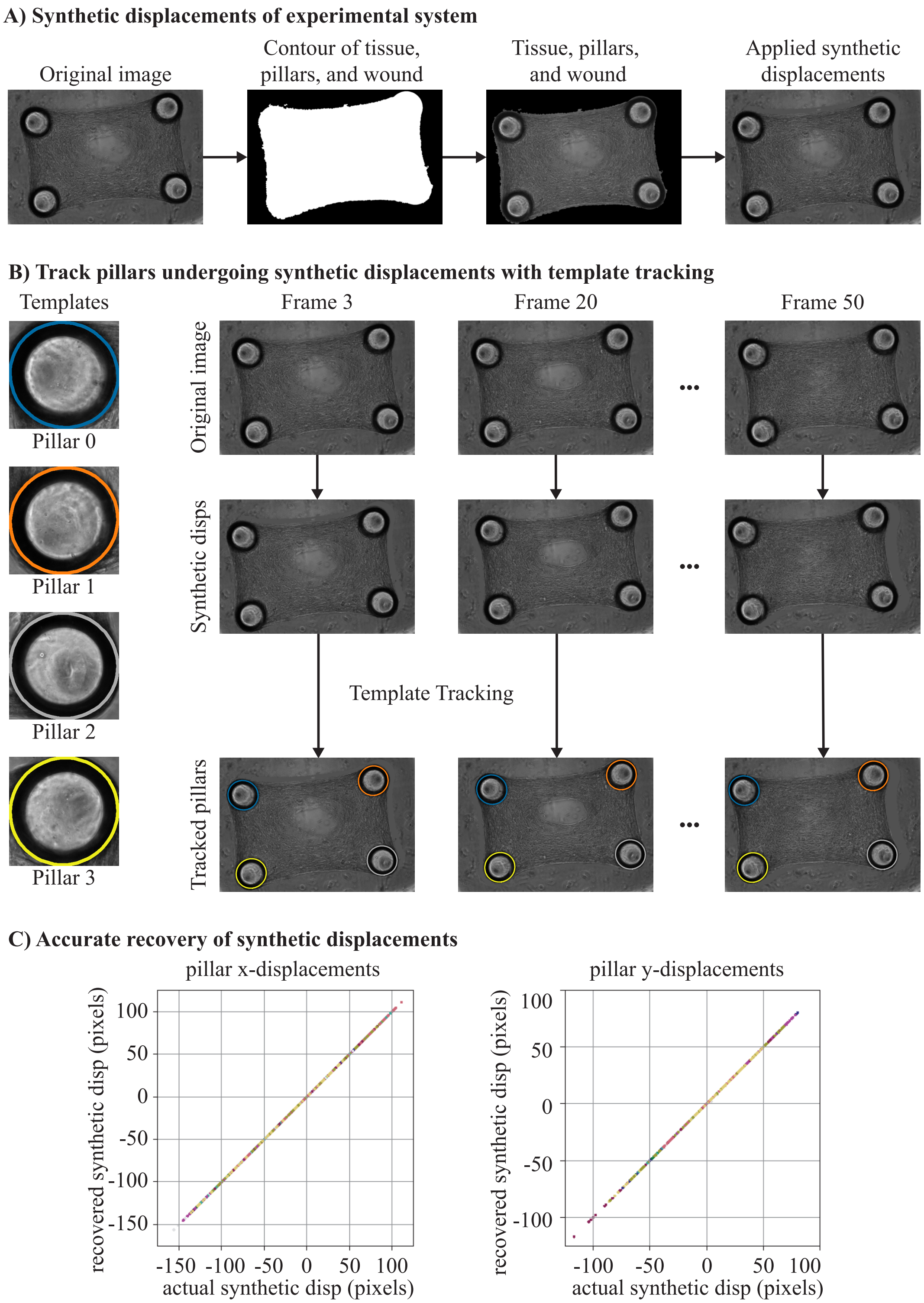


Figure 14: Validation of the pillar tracking algorithm using synthetic displacements. **(A)** Synthetic displacements workflow: the tissue, pillars, and wound are extracted, and displaced by a randomly generated, uniformly distributed synthetic displacement. The displacement magnitude is constrained to keep the sample within the imaging frame. **(B)** Pillar tracking: using templates from the initial frame, the algorithm tracks pillar positions after synthetic displacements. **(C)** Comparison of applied synthetic displacements versus displacements recovered by the tracking algorithm. Further details are provided in Section 4.5.

https://www.ebi.ac.uk/biostudies/bioimages/studies/S-BIAD2719. Additional biological context and experimental motivation are provided in the main manuscript.

Pauli Virtanen, Ralf Gommers, Travis E. Oliphant, Matt Haberland, Tyler Reddy, David Cournapeau, Evgeni Burovski, Pearu Peterson, Warren Weckesser, Jonathan Bright, Stéfan J. van der Walt, Matthew Brett, Joshua Wilson, K. Jarrod Millman, Nikolay Mayorov, Andrew R. J. Nelson, Eric Jones, Robert Kern, Eric Larson, C J Carey, İlhan Polat, Yu Feng, Eric W. Moore, Jake VanderPlas, Denis Laxalde, Josef Perktold, Robert Cimrman, Ian Henriksen, E. A. Quintero, Charles R. Harris, Anne M. Archibald, Antônio H. Ribeiro, Fabian Pedregosa, Paul van Mulbregt, and SciPy 1.0 Contributors. SciPy 1.0: Fundamental Algorithms for Scientific Computing in Python. *Nature Methods*, 17:261–272, 2020. doi:10.1038/s41592-019-0686-2.

Stefan Van der Walt, Johannes L Schönberger, Juan Nunez-Iglesias, François Boulogne, Joshua D Warner, Neil Yager, Emmanuelle Gouillart, and Tony Yu. scikit-image: image processing in python. *PeerJ*, 2:e453, 2014.

Roberto Brunelli. *Template matching techniques in computer vision: theory and practice*. John Wiley & Sons, 2009.

Christopher KI Williams and Carl Edward Rasmussen. *Gaussian processes for machine learning*, volume 2. MIT press Cambridge, MA, 2006.

Saeed Mohammadzadeh and Emma Lejeune. Quantifying hipsc-cm structural organization at scale with deep learning-enhanced sarcgraph. *PLOS Computational Biology*, 21(10):e1013436, 2025.

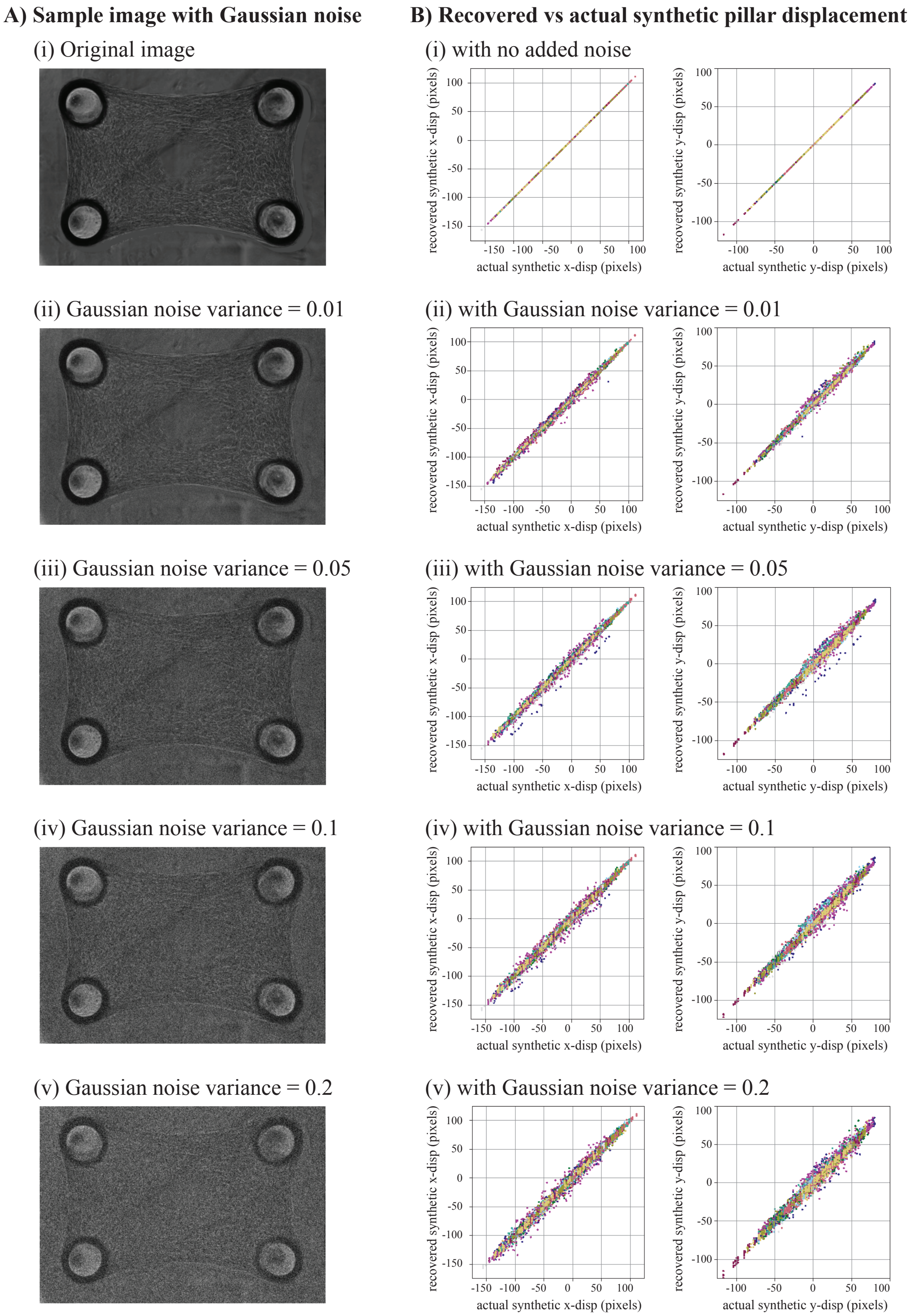


Figure 15: Validation of the pillar tracking algorithm using synthetic displacements with added Gaussian noise.
