## Appendix-S2: Published Dataset Structure for "A high-throughput, 3D microtissue platform for multiparametric analysis of tissue remodeling"

### Appendix S2 - Published Microscopy Dataset

A Preprint

**Quan Nguyen**
Department of Mechanical Engineering, Boston University
Boston, Massachusetts


**Anish Vasan**
Department of Biomedical Engineering and the Biological Design Center, Boston University
Boston, Massachusetts

**Emily Davis**
Department of Biomedical Engineering and the Biological Design Center, Boston University
Boston, Massachusetts

**M Çağatay Karakan**
Department of Biomedical Engineering and the Biological Design Center, Boston University
Boston, Massachusetts

**Elena Westphal**
Department of Biomedical Engineering and the Biological Design Center, Boston University
Boston, Massachusetts

**Varun Shah**
Department of Biomedical Engineering and the Biological Design Center, Boston University
Boston, Massachusetts

**Wilson Wong**
Department of Biomedical Engineering and the Biological Design Center, Boston University
Boston, Massachusetts

**Jeroen Eyckmans**
Department of Biomedical Engineering and the Biological Design Center, Boston University
Wyss Institute for Biologically Inspired Engineering, Harvard University
Boston, Massachusetts

**Emma Lejeune**
Department of Mechanical Engineering, Boston University
Boston, Massachusetts


July 19, 2026

### 1 Introduction

This document describes the published microscopy dataset for our microtissue wound healing experiments, and the settings of our associated software, WoundCompute. We provide metadata and organizational details for the dataset, as well as the input and output data structures required to run WoundCompute on new data.

Reproducibility and reuse of biological imaging data depend critically on the availability of well-organized, well-documented metadata. Without sufficient metadata, it is difficult for other researchers to interpret, reproduce, or build upon published datasets, whether for biological analysis, development of new imaging methods, or training computational models. To address this, we have submitted our dataset to the BioImage Archive (BIA), a free and publicly accessible repository maintained by the European Bioinformatics Institute (EMBL-EBI) to store and distribute biological image data Hartley et al. [2022]. BIA is designed to support FAIR data principles, Findable, Accessible, Interoperable, and Reusable, and requires submitters to provide metadata following the Recommended Metadata for Biological Images (REMBI) guidelines Sarkans et al. [2021]. REMBI organizes metadata into two levels: study-level metadata, which describes parameters that are constant across the dataset as a whole (e.g., imaging modality and microscope settings), and file-level metadata, which describes individual files and their associated experimental conditions (e.g., drug treatment and concentration). Together, these two levels of metadata ensure that our dataset is interpretable by a broad audience, including biologists, imaging scientists, and computational researchers.

This document is organized as follows. Section 2 describes the microscopy dataset, including the experimental setup and the associated REMBI metadata. Section 3 describes the data organization and input and output structures required to run WoundCompute on experimental data. The full dataset is hosted at `https://www.ebi.ac.uk/biostudies/bioimages/studies/S-BIAD2719`.

### 2 Published Dataset

#### 2.1 Microscopy Data Description

Our experimental system comprises 13 96-well plate microtissue wound healing experiments, divided into two groups by cell type: human dermal fibroblasts from adults (NHDFs-Ad) and human dermal fibroblasts from neonatal foreskin (NHDF-Neo). All experiments were conducted using a 96-well plate format, where each well contains a microtissue suspended between four pillars. Laser-ablation was used to injure the microtissues, and time-lapse phase contrast imaging was performed at 30-minute intervals to capture the wound healing response over time (Fig. 1). Experiments differ primarily in cell type and drug condition, with each plate containing one drug treatment alongside one control well. Detailed metadata for all 13 experiments, including frame counts and drug conditions, are provided in Section 2.2.

#### 2.2 Interpreting Associated Metadata

Our dataset is organized into two BIA Study Components, one per cell type, reflecting the two independent experimental cohorts in the dataset. Study-level and file-level metadata are described in the Section 2.2.1 and Section 2.2.2, respectively.

##### 2.2.1 Study-Level Metadata

The study-level metadata describes parameters that are constant across all experiments in the dataset. All images were acquired using a Nikon Eclipse Ti phase contrast microscope at 10$\times$ magnification, with a pixel size of 0.8892 $\mu$m/pixel and a frame interval of 30 minutes. The numerical aperture of the objective is 0.30. The dataset comprises 13 experiments in total, divided into two BIA Biosamples: 11 experiments using NHDFs-Neo and 2 experiments using NHDFs-Ad. Each experiment was conducted using a 96-well plate format, with frame counts ranging from 52 to 98 frames per experiment. Out of 13 experiments, 2 experiments are all-control experiments in which no drug was applied. The remaining 11 experiments each contain one or more drug conditions alongside one control well per plate. A complete summary of all 13 experiments, including cell type, frame count, and drug conditions, is provided in Table 1.

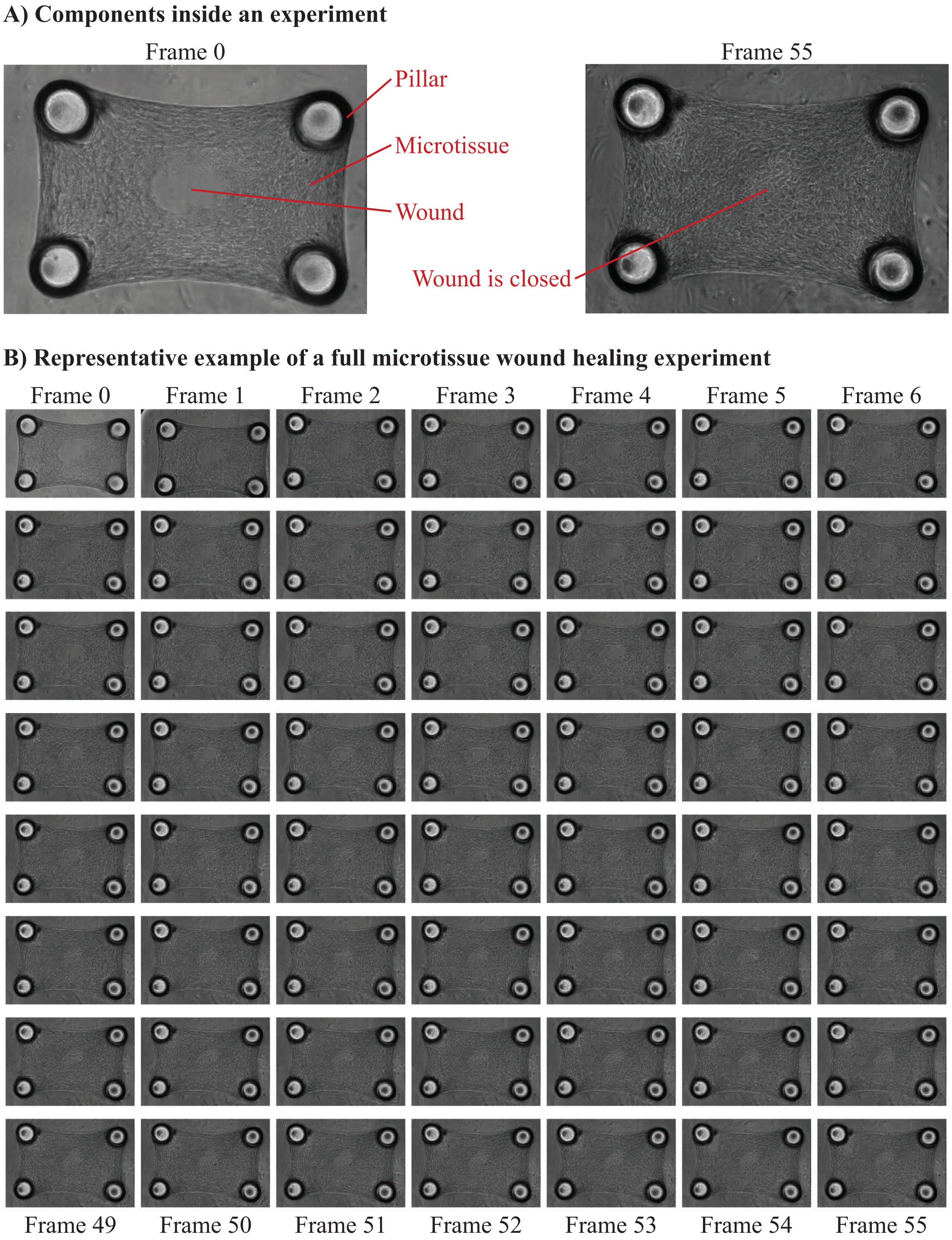


Figure 1: Representative example of a microtissue wound healing experiment. A) Different components in our microtissue experimental system. B) Time-lapse frames from a representative experiment.

Table 1: Summary of all 13 experiments in the dataset with per-well quality control annotations. Each well is annotated to indicate which analyses it was included in after quality control: unmarked wells passed all three (wound area, tissue footprint, and pillar force); superscripts $^{w}$, $^{t}$, and $^{f}$ indicate inclusion in wound area, tissue footprint, or pillar force analysis, respectively (or a subset thereof); † marks wells excluded from all analyses. Conditions shown in *italics* were excluded from all analyses for the reason indicated: [a] bad reagent; [b] seeding or tissue failure; [c] no media change before experiment start; [d] damaged micropillars; [e] no treatment control. All experiments used a 96-well plate format; frame interval is 30 min. Imaging was performed on a Nikon Eclipse Ti phase contrast microscope at 10× magnification (0.8892 $\mu$m/pixel). Full per-well condition and QC assignments are available in `appendix_S2_table_1_well_level_conditions_and_quality_check.xlsx`.

| Experiment | Cell Type | # Fr. | Condition (Concentration) | Wells |
|---|---|---|---|---|
| *BIA Study Component 1: Human dermal fibroblasts from neonatal foreskin (NHDF-Neo)* | | | | |
| R7P1 | NHDF-Neo | 52 | DMSO (0.1%) | A01$^{w}$, A02, A03, A04, A05$^{wt}$, A06†, A07†, A08†, A09$^{tf}$, A10, A11$^{t}$, A12$^{wt}$, B01$^{wf}$, B02†, B03, B04, B05, B06, B07$^{wf}$, B08, B09, B10$^{tf}$, B11$^{w}$, B12, C01$^{wf}$, C02, C03, C04†, C05, C06$^{wt}$, C07$^{t}$, C08$^{wt}$, C09$^{tf}$, C10, C11†, C12$^{wt}$, D01$^{wf}$, D02, D03$^{tf}$, D04†, D05$^{wt}$, D06$^{wt}$, D07†, D08†, D09, D10, D11$^{wt}$, D12$^{w}$, E01$^{tf}$, E02, E03, E04, E05, E06$^{wt}$, E07†, E08, E09, E10, E11$^{w}$, E12†, F01, F02$^{wt}$, F03$^{wf}$, F04, F05$^{wf}$, F06, F07$^{wt}$, F08$^{wt}$, F09†, F10†, F11$^{wt}$, F12†, G01†, G02, G03†, G04†, G05, G06, G07$^{wt}$, G08$^{tf}$, G09$^{tf}$, G10†, G11†, G12$^{w}$, H01, H02$^{wf}$, H03†, H04†, H05, H06$^{wt}$, H07†, H08$^{wt}$, H09$^{wt}$, H10$^{t}$, H11$^{wt}$, H12$^{t}$ |
| R7P2 | NHDF-Neo | 57 | DMSO (0.1%) | A01, A02, A03†, A04$^{w}$, A05†, A06, A07$^{wt}$, A08$^{wt}$, A09†, A10, A11, A12, B01, B02†, B03, B04, B05, B06, B07, B08, B09, B10, B11$^{wf}$, B12, C01$^{wf}$, C02, C03, C04, C05, C06, C07, C08, C09, C10, C11, C12$^{wt}$, D01, D02, D03†, D04†, D05, D06$^{w}$, D07$^{w}$, D08, D09, D10†, D11, D12, E01†, E02, E03, E04, E05, E06, E07, E08$^{wt}$, E09, E10$^{w}$, E11$^{wt}$, E12, F01$^{tf}$, F02, F03$^{wf}$, F04, F05$^{wt}$, F06†, F07†, F08$^{wt}$, F09$^{wt}$, F10, F11$^{wt}$, F12, G01, G02, G03, G04, G05, G06, G07$^{wt}$, G08†, G09†, G10†, G11$^{w}$, G12$^{w}$, H01, H02, H03$^{wt}$, H04, H05, H06, H07$^{w}$, H08$^{wt}$, H09$^{wt}$, H10$^{w}$, H11$^{w}$, H12† |
| R7P5 | NHDF-Neo | 71 | AICAR (20 $\mu$M) | A08$^{t}$, B03$^{wt}$, E01, G08 |
| | | | Blebbistatin (20 $\mu$M) | A11$^{wt}$, D01, E04, G07 |
| | | | Cilengitide (10 $\mu$M) | D05, D11$^{wt}$, G03, G10 |
| | | | Cyclo(RGDyK) (10 $\mu$M) | B05†, D12$^{wt}$, F10, G06$^{w}$ |
| | | | DMSO (0.1%) | A01†, A12, E05, E07 |
| | | | Dinaciclib (5 $\mu$M) | B06, D10, G01, H10 |
| | | | EMD-1214063 (100 nM) | B04$^{wt}$, B09$^{wt}$, F04, G09 |
| | | | Fingolimod (100 nM) | D07$^{wt}$, E12$^{wt}$, F11, H04$^{f}$ |
| | | | Marimastat (5 $\mu$M) | A05$^{tf}$, B12$^{wt}$, E10, F05† |
| | | | Ponesimod (1 $\mu$M) | D06, D09$^{tf}$, E11$^{wt}$, H02 |
| | | | Quercetin Dihydrate (10 $\mu$M) | A09†, D02$^{wt}$, E02, E08 |
| | | | Ruxolitinib (1 $\mu$M) | B10$^{wt}$, D04†, F06, H09 |
| | | | SB431542 (10 $\mu$M) | D08, F12, G11, H05$^{tf}$ |
| | | | Sunitinib (10 $\mu$M) | A04†, B07$^{w}$, F03, F09 |
| | | | Y27632 (10 $\mu$M) | A10$^{wt}$, B02$^{wf}$, E03, H07 |
| | | | *Damaged micropillars*[d] | C01, C03, C08, C09, C10, C11, C12 |
| | | | *Dasatinib (200 nM)*[a] | A07, D03, E09, F01 |

*Continued on next page*

Table 1 – continued from previous page

| Experiment | Cell Type | # Fr. | Condition (Concentration) | Wells |
|---|---|---|---|---|
| | | | *No media change*[c] | G12, H06, H12 |
| R7P6 | NHDF-Neo | 98 | AICAR (20 $\mu$M) | B01, B11, E01, E12[wt] |
| | | | Blebbistatin (20 $\mu$M) | A02†, A07[wt], E04[tf], E07 |
| | | | Cilengitide (10 $\mu$M) | C05, C07, G03, G09 |
| | | | Cyclo(RGDyK) (10 $\mu$M) | C04, C10, G04, G07[tf] |
| | | | DMSO (0.1%) | A01[wt], A12, E05, E06 |
| | | | Dinaciclib (5 $\mu$M) | C06, D01, G02, G10 |
| | | | EMD-1214063 (100 nM) | B07, C01, F03, F10 |
| | | | Fingolimod (100 nM) | D03, G12[w], H06[w] |
| | | | Marimastat (5 $\mu$M) | C03[tf], C12†, G05, G06 |
| | | | Ponesimod (1 $\mu$M) | D02, D12, G01, G11 |
| | | | Dasatinib + Quercetin Dihydrate (200 nM + 10 $\mu$M) | D05, D09[wt], H03, H10 |
| | | | Quercetin Dihydrate (10 $\mu$M) | A05, B12, E10 |
| | | | Ruxolitinib (1 $\mu$M) | B06, C02, F01, F11 |
| | | | SB431542 (10 $\mu$M) | D04, D10[wt], H04, H07 |
| | | | Sunitinib (10 $\mu$M) | B04, B09, F04, F09 |
| | | | Y27632 (10 $\mu$M) | A03†, A06[wt], E03, E09[tf] |
| | | | *Dasatinib (200 nM)*[a] | B02, B10, F05, F06 |
| | | | *No media change*[c] | D06, D07, H01, H11 |
| | | | *Seeding failure*[b] | F02 |
| R7P8 | NHDF-Neo | 87 | Cisplatin (10 $\mu$M) | B11[wt], D07, D12[wt], E02, F01, F10[wt] |
| | | | Crenolanib (10 $\mu$M) | A12[wt], B06, C10[tf], D04, E04[tf], G03[wf], H02 |
| | | | DMSO (0.1%) | A03, B01, C04[wt], F05[wt], G10†, H11[t] |
| | | | Dasatinib (200 nM) | A04[wt], B02, C05, F04, G09[wf], H10 |
| | | | Lenvatinib (10 $\mu$M) | A05, B03†, C06, F03, G08, H06[w] |
| | | | Metformin HCl (10 $\mu$M) | B09, C12, D06, E06, E10, F02, G01 |
| | | | Methotrexate (10 $\mu$M) | D02, D08, E01[wt], E08[tf], E11, E12[wf], F09[wt] |
| | | | Nintedanib (200 nM) | A06[wt], B04, C08, E03[wt], G06[wf], H04[tf] |
| | | | PDGF-BB (100 ng/mL) | C02[w], D01, D09, E09, F08[w], F11†, F12 |
| | | | *Diclofenac (10 $\mu$M)*[a] | A07, B05, C09, D03, G05, H03 |
| | | | *Ibuprofen Lysine (10 $\mu$M)*[a] | B07, B12, D05, D10, E05, G02, H01 |
| | | | *No media change*[c] | A02, C01, C03, E07, F06, G11, G12 |
| R7P9 | NHDF-Neo | 73 | Cisplatin (10 $\mu$M) | C04, C12[wf], D11, E01, E05, F05[tf] |
| | | | Crenolanib (10 $\mu$M) | A10†, B09[wf], D02[tf], E09, F08 |
| | | | DMSO (0.1%) | A02[wt], B02, C09, D05[wt], G07†, H12 |
| | | | Dasatinib (200 nM) | A03†, B03[wf], D06†, D10[tf], G06, H11 |
| | | | Lenvatinib (10 $\mu$M) | A05[w], B04, D07, E10, G05[tf], H10 |
| | | | Metformin HCl (10 $\mu$M) | B12[f], C03[wf], C11†, E07, F01, F06 |
| | | | Methotrexate (10 $\mu$M) | C05[w], D01[wf], D12, E03, E04, F11[wt] |
| | | | *Diclofenac (10 $\mu$M)*[a] | A09, B08, D09, F02, F09, H05 |
| | | | *Ibuprofen Lysine (10 $\mu$M)*[a] | A12, B10, C02, E08, F07, G01 |
| | | | *Nintedanib (10 $\mu$M)*[a] | A06, B05, D08, F10, G02, H08 |
| | | | *No media change*[c] | A01, A04, B01, C08, G08, G12 |
| | | | *PDGF-BB (100 ng/mL)*[a] | C01, C06, D03, D04, F12, G11 |
| R7P10 | NHDF-Neo | 62 | Cisplatin (10 $\mu$M) | B10, D04[wt], D10, D12[wt], E06[tf], F01[tf], F02 |
| | | | Crenolanib (10 $\mu$M) | A10[wt], B07, C08, E09, F03, G05, H02 |
| | | | DMSO (0.1%) | A03[tf], B02, C03, F08, G11, H12† |
| | | | Dasatinib (200 nM) | A04, B03, C04, F07, G10, H07 |
| | | | Lenvatinib (10 $\mu$M) | A05[wt], B04, C05, D07, F06, G09, H06 |
| | | | Metformin HCl (10 $\mu$M) | B09, C10, C12, D03, E07[tf], G01[wt], G02 |

*Continued on next page*

Table 1 – continued from previous page

| Experiment | Cell Type | # Fr. | Condition (Concentration) | Wells |
|---|---|---|---|---|
| | | | Methotrexate (10 $\mu$M) | C11, D05, E01[t], E02, E05, E10, E12 |
| | | | *Diclofenac (10 $\mu$M)*[a] | A09, B06, C07, D09, F04, G07, H03 |
| | | | *Ibuprofen Lysine (10 $\mu$M)*[a] | B08, B12, C09, E03, E08, G03, H01 |
| | | | *Nintedanib (10 $\mu$M)*[a] | A06, B05, C06, D08, F05, G08, H05 |
| | | | *No media change*[c] | A02, C01, C02, F09, F11, G12 |
| | | | *PDGF-BB (100 ng/mL)*[a] | D01, D02, D06, E04, E11, F10, F12 |
| R7P11 | NHDF-Neo | 98 | DMSO (0.1%) | A01, B09, C03, E05† |
| | | | Exendin-4 (100 nM) | A04[wf], C02[wf], C11[wf], D03[wf], F12 |
| | | | Exendin-4 (10 nM) | A03†, B02, D11[wf], H07 |
| | | | Exendin-4 (1 nM) | A02[wf], B03, E02[w], H06[tf] |
| | | | Liraglutide (100 nM) | B07, E06, E12[w], F03[wf], G05[wf] |
| | | | Liraglutide (10 nM) | B08, D01, E07†, F02[tf], G07[wf] |
| | | | Liraglutide (1 nM) | B12, C12, E10[wf], G09 |
| | | | Semaglutide (100 nM) | B01[wt], C01, C06[wf], E09, G11[wt] |
| | | | Semaglutide (10 nM) | A09[wt], C05, D08, D12[t], E11 |
| | | | Semaglutide (1 nM) | A05[wt], C04[wf], D02[wf], E01[f] |
| | | | Tirzepatide (100 nM) | B04[w], E03[wf], F09[wf], F10[tf], H05 |
| | | | Tirzepatide (10 nM) | B05[tf], E04[wf], F07, G02†, G12[tf] |
| | | | Tirzepatide (1 nM) | B06, F01[tf], F06[wf], G04[wf] |
| R7P13 | NHDF-Neo | 91 | DMSO (0.1%) | B12[wt], E04[w], E05, F03[wt], G02[wt] |
| | | | Exendin-4 (100 nM) | A04[wt], B11†, F02[wf], F08[w], H03 |
| | | | Exendin-4 (10 nM) | A03, B07[w], F09, G03†, H04 |
| | | | Exendin-4 (1 nM) | A02, B06[wt], F10, G04[wt], H05[w] |
| | | | Liraglutide (100 nM) | B02[wt], C07, D03, D07, G08, H10 |
| | | | Liraglutide (10 nM) | C06[wf], D01[w], E03, F12[w], G09 |
| | | | Liraglutide (1 nM) | C05, E01, E12, F04[wf], F11[w] |
| | | | Semaglutide (100 nM) | C04, D12[w], E07†, E11[tf], F01, F05 |
| | | | Semaglutide (10 nM) | C02†, C12, D11, F06[wf], G01[w] |
| | | | Semaglutide (1 nM) | A11[w], C11, D02[wf], F07, H02[wt] |
| | | | Tirzepatide (100 nM) | B05[wt], D06, E06, E10[wt], G05[wt], H06[wt] |
| | | | Tirzepatide (10 nM) | B04[wt], D05, D10, G06, H07[wt] |
| | | | Tirzepatide (1 nM) | B03[wf], C10[w], D04[wt], G07, H08 |
| R7P17 | NHDF-Neo | 92 | DMSO (0.1%) | A02, A12[wt], B08, D01[wf], E03, E04, E10, G11[f], G12[wf] |
| | | | Metformin HCl (10 $\mu$M) | A05[w], B04, C08[t], D06, D11, D12[wt], E07, F08[wt], G05, H08† |
| | | | Nintedanib (200 nM) | A11, B07[tf], D08[tf], D10[wt], E05, F03, F05[tf], F11, F12[t], H04 |
| | | | PDGF-BB (100 ng/mL) | A04[wf], B02†, B11[wf], C05, C12, D05[w], E08[wf], F09[wt], G07, G10, H09† |
| | | | PDGF-BB (10 ng/mL) | A03, B10[w], B12[w], C01, C04, D04, E09, F10, H10† |
| | | | *Dasatinib (200 nM)*[a] | A06, B06, C10, D07, E11, E12, F06, G03, H06 |
| | | | *Seeding failure*[b] | B03, D02, G06, H07 |
| EMW15 | NHDF-Neo | 98 | DMSO (0.1%) | A03†, A08†, B04[tf], B08, C01[w], C03†, C07, C12[t], D05, D09†, D10, D11[wf], E06, F02[w], F04, F08†, G01†, G05, G09, G12†, H04, H09 |
| | | | Dasatinib (400 nM) | A06[wt], A10†, B02, B06, B10, C05, C09, D02, D03†, D07, E01†, E04, E08†, E12, F06, F10, F11†, G03, G07, H02, H06[tf], H11 |

*Continued on next page*

Table 1 – continued from previous page

| Experiment | Cell Type | # Fr. | Condition (Concentration) | Wells |
|---|---|---|---|---|
| | | | *Control (untreated)*[e] | A01, A07, A11, B03, B07, C02, C06, C10, C11, D01, D04, D08, E07, F05, F09, F12, G02, G06, G10, H01, H05, H10 |
| | | | *PDGF-BB (10 ng/mL)*[a] | A05, A09, B01, B05, B09, C04, C08, D06, D12, E02, E03, E05, E09, E10, E11, F01, F07, G04, G08, H03, H08, H12 |
| *BIA Study Component 2: Human dermal fibroblasts from adult (NHDF-Ad)* | | | | |
| R7P15 | NHDF-Ad | 90 | DMSO (0.1%) | A01$^{wt}$, C05$^{wt}$, D07, E07, H01$^{wt}$ |
| | | | Exendin-4 (100 nM) | A02$^{wt}$, D01, D08$^{wt}$, E08, F04, H03, H12 |
| | | | Exendin-4 (10 nM) | A03$^{wt}$, B11, D10, E09, F05, H04† |
| | | | Exendin-4 (1 nM) | A04, B12$^{wt}$, D11$^{wt}$, E10, F06, H05 |
| | | | Liraglutide (100 nM) | B04, C10†, E01†, F09$^{wt}$, G03, G11 |
| | | | Liraglutide (10 nM) | B05, C11, D06, F10$^{tf}$, G05, G12 |
| | | | Liraglutide (1 nM) | B06, B07$^{wt}$, D05, F12$^{wt}$, G06, H11 |
| | | | Semaglutide (100 nM) | A05, A11, D12†, E06, E11, H06 |
| | | | Semaglutide (10 nM) | A06†, A10, C08$^{wt}$, E04$^{wt}$, E12, G01 |
| | | | Semaglutide (1 nM) | A09$^{wt}$, B01, C09$^{wt}$, E03$^{wf}$, F08, G02$^{wt}$ |
| | | | Tirzepatide (100 nM) | B08, C01, D04$^{wt}$, F01$^{wt}$, G08, H08, H10$^{wt}$ |
| | | | Tirzepatide (10 nM) | B09, C02$^{wt}$, D03, F02$^{tf}$, G09, H09 |
| | | | Tirzepatide (1 nM) | B10, C03, C06, D02, F03, G10 |
| | | | *Seeding failure*[b] | B03 |
| R7P21 | NHDF-Ad | 86 | DMSO (0.1%) | A01$^{wt}$, A07, B06$^{wf}$, C01$^{w}$, C06$^{w}$, C11†, D06$^{wt}$, E02, E10$^{w}$, E12$^{w}$, F04$^{wt}$, G07$^{w}$, H03$^{wt}$, H09$^{wt}$ |
| | | | Exendin-4 (10 nM) | A02$^{w}$, A08$^{wf}$, B01$^{wt}$, B07, C07†, D02†, D07$^{w}$, D11$^{w}$, F03†, F10$^{wt}$, F12$^{t}$, G06$^{tf}$, H01†, H08 |
| | | | Liraglutide (10 nM) | A04$^{w}$, A10$^{w}$, B03, B09, C03$^{w}$, C09, D03$^{wt}$, D09†, E05$^{wt}$, F01$^{w}$, F08, F11$^{t}$, G04, H06†, H12$^{w}$ |
| | | | PDGF-BB (10 ng/mL) | A06, B05, B11$^{tf}$, C05†, D01$^{w}$, D05, D10$^{wt}$, D12†, E07†, E08$^{w}$, F02$^{wt}$, F05$^{w}$, G10, H04$^{tf}$, H10$^{w}$ |
| | | | Semaglutide (10 nM) | A03$^{t}$, A09, B02, B08$^{tf}$, C02, C08, D08$^{w}$, E03$^{w}$, E04$^{w}$, E11$^{w}$, F09$^{w}$, G01$^{wt}$, G05†, G12$^{wt}$, H07$^{wt}$ |
| | | | Tirzepatide (1 nM) | A05$^{wt}$, B04, B10, C04$^{w}$, C10$^{w}$, C12$^{t}$, D04, E01$^{w}$, E06$^{wt}$, E09$^{wt}$, F06$^{w}$, G03$^{w}$, G11, H05$^{w}$, H11 |
| | | | *Seeding failure*[b] | G08, G09 |

##### 2.2.2 File-Level Metadata

File-level metadata is provided in three spreadsheets submitted alongside the dataset to BIA, one per file category. These spreadsheets store the necessary metadata for users to understand what each file contains and how it relates to other files in the dataset. The three spreadsheets, containing raw experimental images, segmentation masks, and pillar tracking data, are described below.

**Raw Images – `metadata_experimental_images.xlsx`** This spreadsheet contains one row per raw `.TIF` image file. The columns are described in Table 2.

**Segmentation Masks – `metadata_mask_annotations.xlsx`** This spreadsheet contains one row per segmentation mask file. The columns are described in Table 3.

**Pillar Tracking Data – `metadata_pillar_position_annotations.xlsx`** This spreadsheet contains one row per pillar tracking output file, encoding a temporal relationship: the source fields describe the reference frame from which pillar templates are extracted, while the output fields describe files spanning the full time series of a given sample. The columns are described in Table 4.

Table 2: Column descriptions for `metadata_experimental_images.xlsx`.

| Column Name | Description |
|---|---|
| `File` | Path to the raw `.TIF` image file |
| `plate_id` | 96-well plate identifier, corresponding to experiment names in Table 1 |
| `well_number` | Well position within the plate |
| `condition` | Experimental condition applied to the well |
| `condition_concentration` | Concentration of the drug applied |
| `time_between_frames_in_mins` | Imaging frame interval in minutes (30 for all entries) |

Table 3: Column descriptions for `metadata_mask_annotations.xlsx`.

| Column | Description |
|---|---|
| `File` | Path to the segmentation mask `.npy` file |
| `source_image` | Path to the raw `.TIF` image from which the mask was segmented; each mask corresponds to exactly one source image |
| `annotated_object` | Segmented object of interest: `pillar`, `tissue_footprint`, or `wound` |

### 3 Running WoundCompute on Experimental Data

This Section describes the data organization and file structures required to run WoundCompute on experimental data. We cover three stages: the raw data as exported from the microscope, the organized raw data prepared for WoundCompute input, and the analysis outputs produced by WoundCompute. Figure 2 illustrates the full data organization. For the software to function without modification, it is required to follow the file structure organization described here.

#### 3.1 File Naming Conventions

Raw data exported from the microscope follows the naming convention: [tissue status information]_s[sample index]_t[time index].TIF (Fig. 2A). For example, `tissue_ai_s01_t0001.TIF` corresponds to a tissue after injury, sample 1, and frame 1. This systematic naming convention allows the data to be organized automatically, as described in Section 3.2.

#### 3.2 WoundCompute Input – Organized Raw Data

Before running WoundCompute, the raw data must be organized as shown in Fig. 2B, with `.TIF` files sorted by experiment and sample index. This structure ensures that the software processes all frames from the same sample in a single run. The accompanying `.yaml` file defines the analysis parameters (details in Section 3.3). For convenience, the WoundCompute GUI (available at `https://github.com/quan4444/woundcomputeGUI`) can automatically format the data and generate the necessary `.yaml` files. The `organize_files` function (`https://github.com/quan4444/woundcomputeGUI/blob/main/src/woundcomputegui/main_gui.py`) provides the underlying code for data organization.

#### 3.3 WoundCompute Settings – Input `.yaml` File

Since WoundCompute supports multiple analysis types and imaging modalities, we use a `.yaml` file to specify which analyses to run and to document settings for future reproducibility. Here is an example `.yaml` file:

```yaml
# User input for running the test_movie example
version: 1.0 # do not modify
segment_brightfield: false
seg_bf_version: 1 # do not modify
seg_bf_visualize: false
segment_fluorescent: false
seg_fl_version: 1 # do not modify
seg_fl_visualize: false
segment_ph1: true
seg_ph1_version: 2
seg_ph1_visualize: true
track_brightfield: false # do not modify
track_bf_version: 1 # do not modify
```

Table 4: Column descriptions for `metadata_pillar_position_annotations.xlsx`.

| Column | Description |
|---|---|
| `File` | Path to the pillar position `.txt` file |
| `source_image` | Path to frame 0 of the sample, used as the reference frame for tracking |
| `source_pillar0_mask` | Path to the binary mask `.npy` file for pillar 0, extracted from the reference frame |
| `source_pillar1_mask` | Path to the binary mask `.npy` file for pillar 1, extracted from the reference frame |
| `source_pillar2_mask` | Path to the binary mask `.npy` file for pillar 2, extracted from the reference frame |
| `source_pillar3_mask` | Path to the binary mask `.npy` file for pillar 3, extracted from the reference frame |
| `tracked_tif_files` | File names of all frames across which pillar positions were tracked |
| `x_or_y_position` | Denotes whether the file contains $x$- or $y$-position data |

```
track_bf_visualize: false # do not modify
track_ph1: false # do not modify
track_ph1_version: 1 # do not modify
track_ph1_visualize: false # do not modify
bf_seg_with_fl_seg_visualize: false
bf_track_with_fl_seg_visualize: false # do not modify
ph1_seg_with_fl_seg_visualize: false # do not modify
ph1_track_with_fl_seg_visualize: false # do not modify
zoom_type: 2 # set to 1 for examples where you cannot see the pillars,
             # set to 2 for examples where the pillars are visible
track_pillars_ph1: true # set to False for zoom type 1,
                        # set to True for pillar tracking with zoom type 2
segment_dic: false
seg_dic_version: 1 # do not modify
seg_dic_visualize: false
track_dic_visualize: false
track_pillars_dic: false
run_before_injury_and_after_injury_together: false
low_quality_frame_inds: []
```

Fields marked `# do not modify` correspond to features that have been implemented but not yet thoroughly tested and validated. These features are reserved for future work and should not be modified by the user at this time.

For running WoundCompute on phase contrast data, the `.yaml` file should look identical to the example above. Any analysis step can be skipped by toggling its corresponding field to `false`. For example, if your experiment does not include fluorescent imaging, set `segment_fluorescent`, `seg_fl_visualize`, and `bf_seg_with_fl_seg_visualize` to `false` to skip fluorescent segmentation and visualization entirely. The `low_quality_frame_inds` field accepts a Python-style list of integer frame indices (zero-indexed) to exclude from use as reference frames for pillar template extraction and tracking; for example, `low_quality_frame_inds: [0, 3, 7]` excludes the first, fourth, and eighth frames. If no frames need to be excluded, this field should be left as an empty list: `low_quality_frame_inds: []`. Note that frames listed here are still processed through the full analysis pipeline.

##### 3.4 WoundCompute Output File Structure

After WoundCompute analysis, output files are written to the `/segment_ph1` and `/track_pillar_ph1` folders inside each sample folder, as illustrated in Fig. 2C. All mask files are 2D boolean arrays with shape $(H, W)$, where $H$ and $W$ match the dimensions of the input images and are consistent across all frames within a sample and across all samples within an experiment. Wound area and axis length values are reported in pixels; to convert to physical units, multiply by the pixel size of 0.8892 $\mu$m/pixel. The output files are as follows:

- `tissue_footprint_mask_[time index].npy`: 2D boolean array of shape $(H, W)$ representing the tissue footprint mask at each time step.
- `wound_mask_[time index].npy`: 2D boolean array of shape $(H, W)$ representing the wound mask at each time step.
- `contour_coords_[time index].npy`: array of shape $(N, 2)$ containing the coordinates of the wound contour at each time step, where $N$ is the number of contour points and may vary across frames. Values are stored as floats.

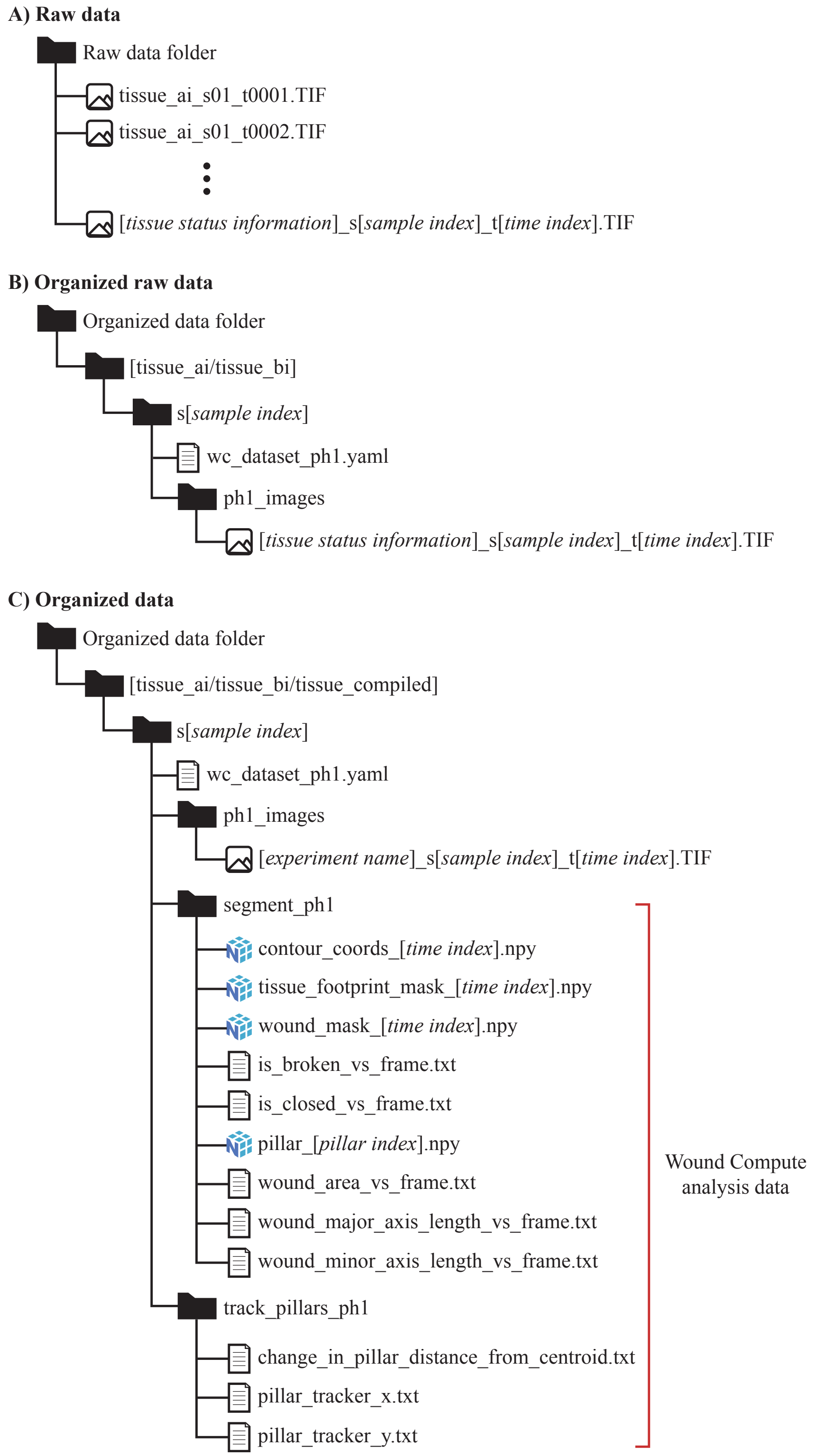


Figure 2: Dataset organization: A) Raw data from microscopy imaging experiments, B) Organized raw data, C) All organized data including the WoundCompute analysis outputs.

- `pillar_[pillar index].npy`: 2D boolean array of shape $(H, W)$ representing the binary mask of each pillar, extracted from the reference frame.
- `is_broken_vs_frame.txt`: tissue broken status over time. Contains one binary value per line, where 0 indicates the tissue is intact and 1 indicates the tissue is broken. See Appendix S1 (WoundCompute Software) for details on how broken status is determined.
- `is_closed_vs_frame.txt`: wound closure status over time. Contains one binary value per line, where 0 indicates the wound is open and 1 indicates the wound is closed.
- `wound_area_vs_frame.txt`: wound area over time. Contains one value per line, in pixels.
- `wound_area_vs_frame_GPR.txt`: wound area over time, smoothed using Gaussian Process Regression (GPR) with a Radial Basis Function (RBF) kernel. Contains one value per line, in pixels.
- `wound_area.png`: plot showing both the raw and GPR-smoothed wound area over time.
- `wound_major_axis_length_vs_frame.txt`: length of the major axis of the wound at each time step. Contains one value per line, in pixels.
- `wound_minor_axis_length_vs_frame.txt`: length of the minor axis of the wound at each time step. Contains one value per line, in pixels.
- `pillar_positions.png`: image displaying the assignment of pillar indices to their corresponding pillars in the field of view.
- `pillar_tracker_x.txt`: $x$-positions of all 4 pillars over time. Each line corresponds to one frame, with pillar positions separated by spaces.
- `pillar_tracker_y.txt`: $y$-positions of all 4 pillars over time. Each line corresponds to one frame, with pillar positions separated by spaces.
- `change_in_pillar_distance_from_centroid_[sample name].png`: plot showing the change in distance of each pillar from the centroid of all pillars over time.
- `change_in_pillar_distance_from_centroid.txt`: change in distance of each pillar from the centroid of all pillars over time. Each line corresponds to one frame, with one value per pillar separated by spaces. For example, a sample with 4 pillars and 52 frames will have 52 lines, each containing 4 values.

The algorithms and computational procedures underlying each of these outputs are described in detail in Appendix S1 (WoundCompute Software).

#### 4 Data and Code Availability

The full dataset is publicly available through the BioImage Archive at `https://www.ebi.ac.uk/biostudies/bioimages/studies/S-BIAD2719`. WoundCompute source code is available at `https://github.com/elejeune11/woundcompute` (archived on Zenodo, `https://doi.org/10.5281/zenodo.19830254`), and a graphical interface for non-computational users is available at `https://github.com/quan4444/woundcomputeGUI`. Further documentation for both is provided in the respective README files and in Appendix S1 (WoundCompute Software). Additional biological context and experimental motivation are provided in the main manuscript.
